# Anaerobic riboflavin degradation by human gut *Lachnospiraceae*

**DOI:** 10.64898/2026.02.13.705874

**Authors:** Christian J. Quiles-Pérez, Alexandra Olzak, Aminata Fofana, Kamal Deep, Calissa Carlisle, Emily Bradley, Kathryn Kananen, Connor Skaggs, Justin A. North, Patrick H. Bradley

## Abstract

Vitamins mediate a web of cross-feeding interactions in the human gut. Many Gram-positive gut microbes, in particular, are predicted to be vitamin auxotrophs. Previous studies of these microbes, however, have tended to use rich media, precluding controlled perturbations of low abundance nutrients. We tested the ability of diverse *Lachnospiraceae*, the most common Gram-positives in the gut, to grow on a chemically defined medium. Even though this medium contained riboflavin, we found that predicted riboflavin auxotrophs grew poorly, including the bile metabolizer *Clostridium scindens*. High-dose riboflavin supplementation enhanced growth, but also revealed that surprisingly, *C. scindens* catabolizes riboflavin into lumichrome, making it the first reported anaerobe to do so. The only previously described catabolic pathway for riboflavin requires oxygen and has no homologs in *C. scindens*. In high-dose riboflavin, a single gene neighborhood with an aldolase, oxidoreductases, and a riboflavin kinase/adenylyltransferase was upregulated, suggesting an alternative anaerobic degradation or overflow pathway. Similar neighborhoods were detected in several other *Lachnospiraceae*, including *Faecalicatena fissicatena*, the only other anaerobe reported to degrade riboflavin. Reanalysis of published metabolomic data showed that *in vivo*, both riboflavin and lumichrome were more abundant in colonized (vs. germ-free) mouse ceca, and that *in vitro, Lachnospiraceae* isolates depleted riboflavin while certain Gram-negative isolates overproduced it. These results demonstrate that a member of the *Lachnospiraceae* can anaerobically convert an essential B vitamin into lumichrome, a molecule recently shown to have anti-inflammatory properties. Vitamin catabolism may both structure cross-feeding interactions in the gut and affect host health.

**Importance:** *Lachnospiraceae*, the most prevalent Gram-positives in the human gut, produce many health-relevant metabolites, but are genetically intractable and are often grown in rich medium, complicating physiological studies. By performing a comparative study with chemically defined media, we identify for the first time a specific anaerobe that can break down riboflavin to lumichrome. Using transcriptomics, we also identify a specific gene neighborhood representing the first candidate pathway for anaerobic flavin degradation. This neighborhood is conserved in a handful of other *Lachnospiraceae*, including *F. fissicatena*, the only other anaerobe known to degrade riboflavin (to the related product hydroxyethylflavin). These results may explain decades-old observations implicating gut microbes in the formation of riboflavin degradation products. Furthermore, lumichrome and related metabolites have recently been shown to inhibit host mucosal-associated invariant T (MAIT) cell activation, suggesting an additional mechanism by which commensal *Lachnospiraceae* may dampen inflammation.

## Introduction

Gut microbial metabolism drives many of its links to host health: certain microbial metabolites can feed host cells^1^, regulate immune responses^2^, and inhibit pathogens^3,4^, while others may, for example, worsen renal disease^5^ or predispose the host to atherosclerosis^6,7^. Modulating this metabolic activity therefore offers a route towards novel gut-targeted therapeutics, whether via small molecule inhibitors of microbial pathways^8^ or altering the nutritional environment, e.g., by providing prebiotics^9^. Prebiotics modulate the dietary molecules available to gut microbes; however, the gut microbiome also extensively metabolizes host molecules (e.g., mucin, urea, lactate^10–12^) as well as nutrients from other members of the community, via cross-feeding^13^.

Cross-feeding between members of the gut microbiome may be particularly relevant for vitamins^13^. Vitamins are usually required in small amounts, and the host efficiently absorbs many vitamins in the proximal small intestine^14^. Thus, in the absence of large-dose supplementation that overwhelms this machinery, microbial sources of vitamins may be especially important *in vivo*. In synthetic communities where auxotrophies of various types were introduced by gene deletion, vitamin auxotrophy specifically was found to impose low costs on growth rate while leading to stable cross-feeding relationships^15^. Many gut microbes are predicted based on genome sequences to be vitamin auxotrophs^16^, and co-culture experiments have confirmed that auxotrophs for quinones^17^ or B vitamins^18^ can indeed be fed by gut prototrophs.

Vitamin availability may also affect which specific pathways gut microbes can perform, and therefore which specific products are formed. For example, pseudovitamin B_12_ provision by *E. hallii* shifts carbon metabolism in the mucus degrader *A. muciniphila* to favor the propionate over succinate^19^, while supplementation with very high-dose riboflavin has been shown to shift gut community metabolism towards the production of butyrate^20^ without major alterations to community structure. Both propionate and butyrate are health-associated short-chain fatty acids (SCFAs) that modulate a range of host phenotypes ranging from immunity to metabolism^21^, illustrating the potential impact of vitamin-dependent metabolic remodeling.

As redox balance is one of the most important determinants of metabolic routes in gut anaerobes^22^, vitamins that form redox-active cofactors, like nicotinate and flavins, are good candidates for influencing gut microbial metabolism. Flavins play especially critical roles for anaerobes. Flavodoxins often substitute for ferredoxins in iron-limited environments, and are therefore critical for activating certain radical enzymes as well as providing electrons to hydrogenase. (Hydrogen itself is an important redox “currency metabolite” in the gut, whose concentration can itself affect metabolic flux^23^.) Flavoproteins also transfer electrons to alternative acceptors, such as urocanate and fumarate^24^. Yet another possible role for flavins is serving as an extracellular electron shuttle, as demonstrated in *Shewanella*^25^, *Listeria*^26^, and potentially the gut commensal *Faecalibacterium prausnitzii*^27^ (a member of the Clostridia, though not the *Lachnospiraceae*), which has been reported to reduce soluble riboflavin to reduce oxygen “at a distance,” similarly to how *P. aeruginosa* in biofilms are thought to use the structurally-similar phenazines^28,29^.

With some exceptions, vitamins are thought to be mainly long-lived molecules that are extensively recycled, explaining why they are typically required only at low concentrations. However, vitamins may also be transformed by microbes. One type of transformation involves converting vitamins into forms that may be more or less accessible by particular taxa, as seen with, for example, vitamin B _12_ ^30,31^. Cofactors containing vitamins may also be susceptible to metabolic damage through side or spontaneous reactions^32^. Perhaps counterintuitively, microbes may also actively degrade vitamins, and potentially even use them as additional carbon sources if they are available in high enough concentrations, as seen with nicotinate fermentation in *E. barkeri*^33^, and riboflavin catabolism in *M. maritypicum*^34^ and *D. riboflavina*^35^. Breakdown products of riboflavin have also previously observed in the mammalian gut and in urine, with gut microbes implicated in their production^36–38^.

Despite this, the role of vitamin degradation or catabolism in the gut microbial food web has received relatively little attention. One potential reason is that the mechanisms for such degradative pathways have not yet been elucidated, making it difficult to predict this activity from (meta)genomics data. The aerobic riboflavin degradation pathway, for instance, was only worked out in the late 2010s^34,35^ even though this activity was initially observed as early as 1944^39^. Furthermore, as a key step of this pathway requires oxygen^34,35^, there is currently no explanation for how anaerobic microbes might perform a similar activity.

In the present study, we focus on the *Lachnospiraceae*, which are the most common Gram-positive family in the human gut^40^ and are predicted to be auxotrophic for several B vitamins^16,18^. As these microbes produce a variety of health-relevant molecules^3,41–43^, we were motivated to investigate what growth factors they require, and how nutrient availability may regulate the production of these key metabolic products. Chemically defined media are powerful tools for answering questions like these^44^, as they allow precise perturbations of even low-abundance nutrients. However, while a chemically defined medium has been developed for at least one species of the *Lachnospiraceae*^41^, and a semi-defined medium has been developed for certain others^18^, most isolates are still typically grown in rich media. *Lachnospiraceae* also unfortunately remain largely genetically intractable, further limiting our ability to investigate metabolic regulation. To address these limitations, we therefore took a comparative approach^45^, using natural variation to link differences in growth to genetic variation. Furthermore, our experiments use a fully chemically-defined medium, which we modeled on previous successful work in *C. scindens*^41^ and other *Lachnospiraceae*^18^.

## Results

### The ability to synthesize riboflavin *de novo* explains variation in *Lachnospiraceae* growth on a chemically defined medium (DM)

We grew five diverse *Lachnospiraceae* in two variants of a chemically defined medium: one had fructose as a sole carbon source (DM), and one was additionally supplemented with acetate (DM+Ace). To test for sustained growth, we measured optical density across three successive subcultures. *Robinsoniella peoriensis, Anaerostipes caccae, and Dorea formicigenerans* consistently exhibited robust growth on DM, reaching a stable maximum OD similar to what they reached on the rich medium BHI+CHK (Brain Heart Infusion with L-cysteine, hemin, and Vitamin K; Figure 1A-B). In contrast, *Clostridium scindens* and *Blautia coccoides* had growth defects, with *C. scindens* growing a very small amount in each subculture (maximum OD near 0.1), and *B. coccoides* showing no growth at all past the first subculture. Adding acetate slightly increased the growth rate of *D. formicigenerans* and *C. scindens*, but without changing the pattern we observed in maximum OD across species.

**Figure 1:**
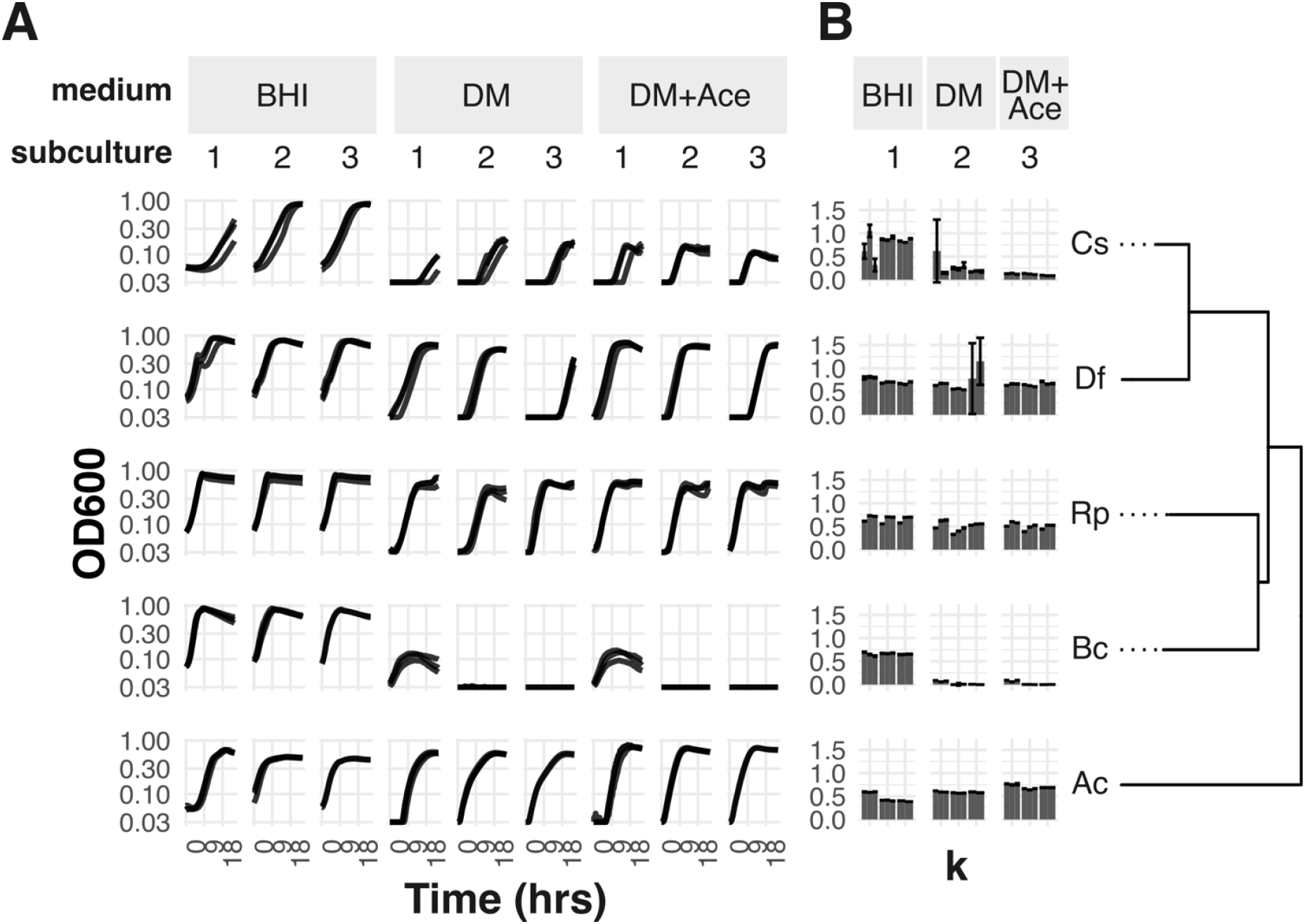
Growth of Lachnospiraceae isolates in rich and defined media. A) Growth curves in Brain Heart Infusion (BHI, biological n=4), defined medium with fructose as the carbon source (DM, n=3), and defined medium with both fructose and acetate (DM+Ace, n=3). Each line corresponds to one biological replicate and is the average of 4 technical replicates. Subcultures are 20-fold dilutions into fresh media. B) Estimated carrying capacity (k) for each biological replicate (individual bars) based on a logistic curve fit. Error bars are 2xSE; one data point with an SE>0.5 was dropped. Species abbreviations are as follows: Cs, *Clostridium scindens*; Df, *Dorea formicigenerans*; Rp, *Robinsoniella peoriensis*, Bc, *Blautia coccoides*, Ac, *Anaerostipes caccae*.

We set out to explain these differences based on gene content. This required us to first sequence the *R. peoriensis* type strain, which at that time did not have a full genome sequence. Using a combination of long- and short-read sequencing (see Methods), we were able to obtain a genome with only two scaffolds. Next, we annotated genomes for all five species using a consistent pipeline based on anvi’o, which we previously found had higher sensitivity to detect KEGG Ortholog families (KOfams) in taxa like the *Lachnospiraceae*^46^. We found that only 10 KOfams were present in *R. peoriensis, A. caccae*, and *D. formicigenerans* but not *C. scindens* and *B. coccoides* (Figure 2A), and of these, five belonged to a single pathway: *de novo* riboflavin biosynthesis (Figure 2B; full set in Supplementary Table S2 and Supplementary Table S3).

**Figure 2:**
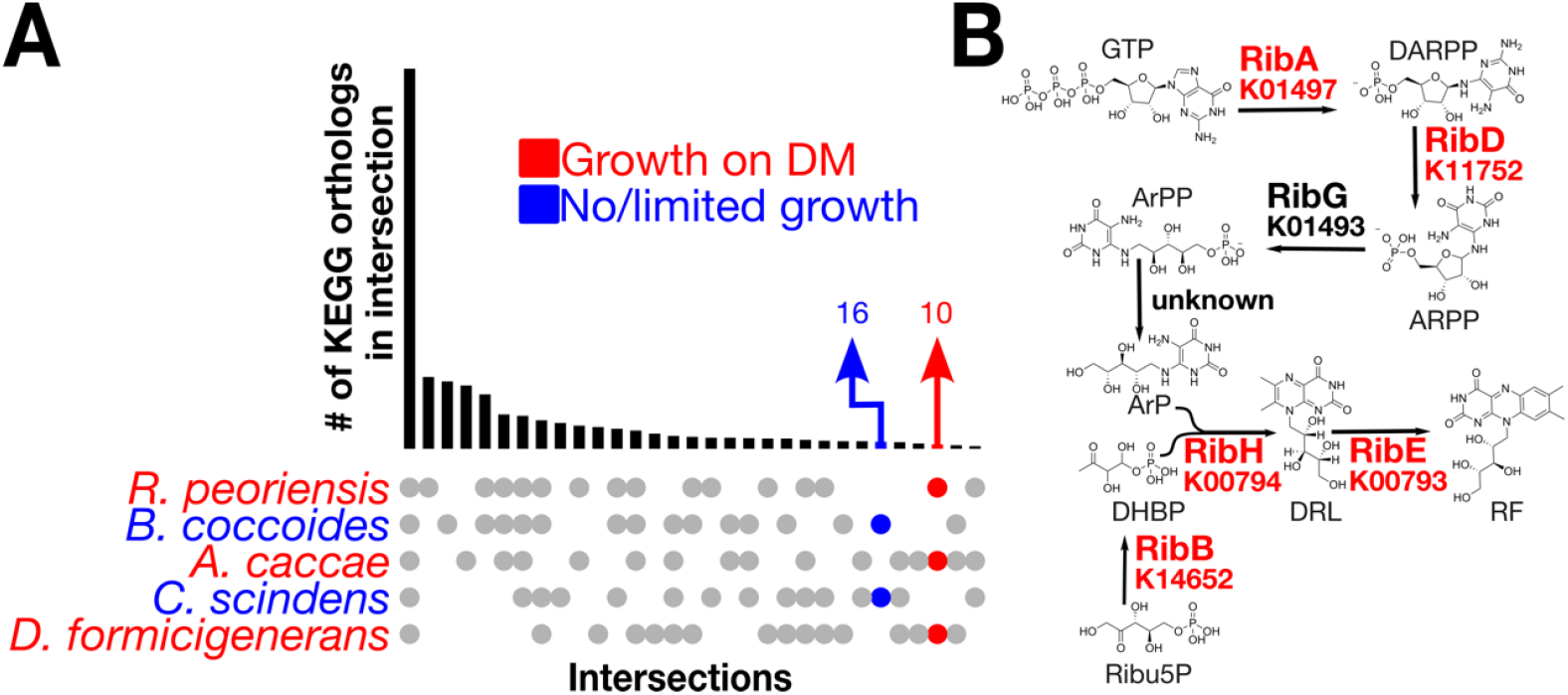
The de novo riboflavin biosynthesis pathway explains poor growth on DM. A) UpSet plot of KEGG Orthologs showing the overlap (bars) for a given intersection of genomes (filled circles). The intersection of taxa that did not grow well is colored blue, while the intersection that did grow is colored red. B) Diagram showing the *de novo* riboflavin biosynthesis pathway. Genes present in the strains that grew well are highlighted in red. DARPP: 2,5-diamino-6-(1-D-ribosylamino)pyrimidinone phosphate; ARPP: 5-amino-6-(1-D-ribosylamino)pyrimidinone phosphate; ArPP: 5-amino-6-(1-D-ribitylamino)pyrimidinone phosphate; ArP: 5-amino-6-(1-D-ribitylamino)pyrimidinone; Ribu5P: ribulose-5-phosphate; DHBP: 3,4-dihydroxy-2-butanone 4-phosphate; DRL: 6,7-dimethyl-8-ribityllumazine; RF: Riboflavin.

These results pointed to riboflavin auxotrophy as a potential explanation; indeed, *C. scindens* ATCC 35704 is already known to require riboflavin to grow^41^. However, the DM we used did contain riboflavin at 0.13µM, a concentration that was previously shown to permit growth for three other riboflavin auxotrophs from the gut (e.g. *R. inulinivorans*)^18^.

Because flavins play such important roles in anaerobic metabolism, one possibility is that *C. scindens* might simply need much higher quantities of riboflavin. As mentioned, flavodoxin plays an important role in activating radical enzymes and in providing high-energy electrons to hydrogenase; *C. scindens* is a prolific producer of hydrogen^41^, and both *C. scindens* and *B. coccoides* encode hydrogenase maturation cofactors absent from the other three strains we tested (Supplementary Table S2 and Supplementary Table S3). *C. scindens*, in particular, is also known to use flavoproteins to reduce bile acids and steroids^47,48^. Finally, flavins (including, unusually, riboflavin itself), are also components of Rnf/Nqr-like systems^49^, which are present in *C. scindens*. If *C. scindens* and/or *B. coccoides* simply require larger amounts of riboflavin to grow well, then riboflavin auxotrophy could still explain the pattern of growth we observed.

### Riboflavin supplementation strongly enhances *C. scindens* growth

We first tested the hypothesis that the concentration of riboflavin we used in our DM was limiting for *C. scindens*. Indeed, when supplemented with 17.5µM riboflavin, *C. scindens* showed a more than two-fold increase in peak OD (Figure 3A). Riboflavin supplementation did not affect the growth of *B. coccoides* or *R. peoriensis*.

**Figure 3:**
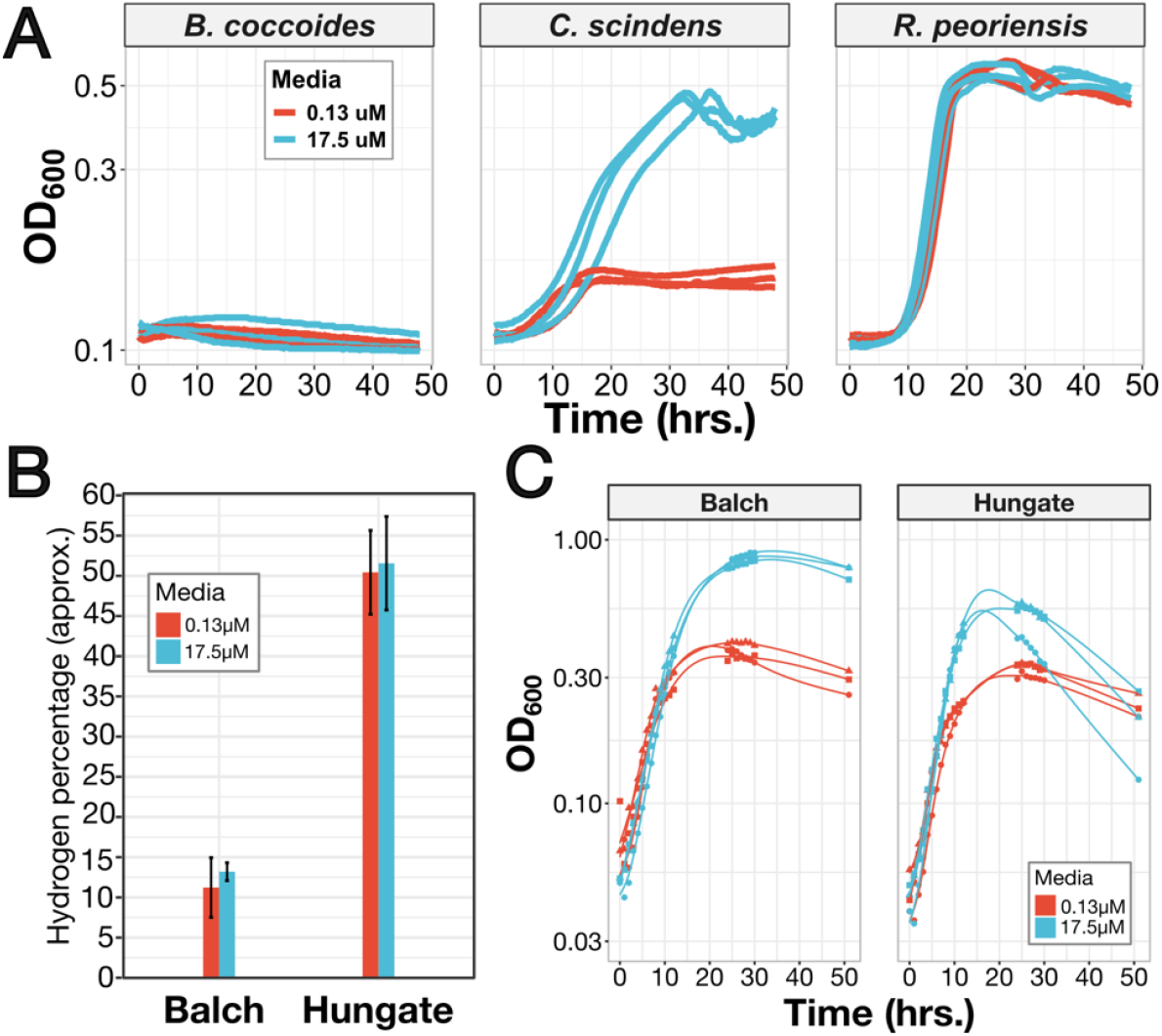
High-dose riboflavin enhances growth of *C. scindens*, independently of relief of hydrogen inhibition. A) Growth curves of *B. coccoides, C. scindens*, and *R. peoriensis* on DM in the presence of either 0.13µM (red) or 17.5µM (teal) riboflavin. Each line is one biological replicate (n=3 total) and is the average of n=2 technical replicates. B) Hydrogen concentrations in Balch and Hungate tubes at 51 hrs. growth (n=3 biological replicates; error bars are 1xSE), estimated from gas chromatography. Note that percentages are estimates because the very high H_2_ concentrations were all above the concentrations we used in our calibration curve. C) Growth curves of C. scindens grown on 0.13µM (red) and 17.5µM riboflavin (teal), in either Balch tubes with large headspaces (left) or Hungate tubes with small headspaces (right).

One potential reason we did not see strong growth with 0.17µM riboflavin could be low or inefficient transport across the membrane. This is observed in, for example, *E. coli* riboflavin auxotrophs, which lack a dedicated transporter^50,51^ and require hundreds of micromolar riboflavin to grow. If a transporter is heterologously expressed in those strains, they then require only micromolar amounts to grow to a high OD^52^. However, we found genes encoding homologs of the RibU riboflavin transporter^53^ (KOfam K23675) in all five strains that we tested, and actually observed a second RibU paralog in both *B. coccoides* and *C. scindens*. In *L. lactis*, where RibU was discovered, the *ribU* gene is regulated by an upstream FMN riboswitch^54^ (Rfam RF00050), a gene regulatory element that acts as a terminator when bound to intracellular FMN^55^. Consistent with a role in riboflavin transport, RefSeq^56^ annotations for both *C. scindens* and *B. coccoides* also included an FMN riboswitch element upstream of each copy of *ribU*. This strongly suggests the presence of functional riboflavin transporters.

### Riboflavin does not enhance *C. scindens* growth by relieving hydrogen inhibition

Even with supplementation to 17.5µM, the maximum density of *C. scindens* we observed was still much lower than what was obtained previously on a very similar defined medium containing 1.7µM riboflavin, which was actually designed for this exact strain^41^. One potential explanation was that we initially cultured *C. scindens* in 96-well plates inside an anaerobic chamber, in which the gas atmosphere contained 10% CO_2_ and between 2–3% H_2_. In contrast, the experiments where *C. scindens* grew best used glass Balch tubes containing a defined gas atmosphere of 100% CO_2_ ^38^. *C. scindens* is known to produce hydrogen itself^41^, and hydrogenase can be easily product-inhibited. Furthermore, while hydrogen has low solubility in water, cultures in a 96-well plate would have much higher surface-to-volume ratios than Balch tube cultures, and so we would expect more exposure to atmospheric H_2_ during plate growth inside the chamber. Hydrogenase inhibition could cause reduced flavins to accumulate without a way to re-oxidize them (for example, because flavodoxin is the electron donor to hydrogenase in low iron conditions); if this were true, riboflavin supplementation could enhance growth by providing a source of oxidized flavins. We therefore set out to test whether riboflavin still enhanced *C. scindens* growth when hydrogen product inhibition was relieved.

To do so, we grew *C. scindens* in our DM with 0.13µM and 17.5µM riboflavin in glass anaerobic culture tubes without shaking, using a 100% CO_2_ atmosphere (see Methods). Furthermore, we used a consistent 10mL culture volume in two different sizes of tubes, Hungate tubes and Balch tubes. This allowed us to control the headspace, which was approximately 3mL in Hungate tubes and 17mL in Balch tubes. As expected, hydrogen accumulated to a much higher concentration in the headspace of the Hungate tubes than the Balch tubes (Figure 3B). Regardless of the amount of riboflavin provided, the final OD was significantly higher in the Balch tubes, consistent with less inhibition of hydrogenase due to the lower H_2_ concentrations; we also noted that Hungate tube cultures showed a drop in OD after hitting the maximum, which could be a result of cell death or a stress-induced change in cell morphology. However, riboflavin supplementation increased the peak OD by almost the same factor in both Hungate and Balch tubes (Figure 3C). Therefore, we conclude that hydrogen (and likely, hydrogenase inhibition) can explain why we saw poorer growth of *C. scindens* overall, but also, that the mechanism by which riboflavin enhances *C. scindens* growth must be independent.

### *C. scindens* breaks down riboflavin into lumichrome anaerobically

An alternative explanation for *C. scindens’* high need for riboflavin could be that flavins were being degraded during growth, requiring replenishment beyond what would be necessary to account for dilution. In support of this hypothesis, we noticed that while the 17.5µM riboflavin medium was bright yellow, *C. scindens* cultures turned the medium clear over the course of six hours. Uninoculated controls remained brightly colored over the same time period (Figure 4B, insets).

**Figure 4:**
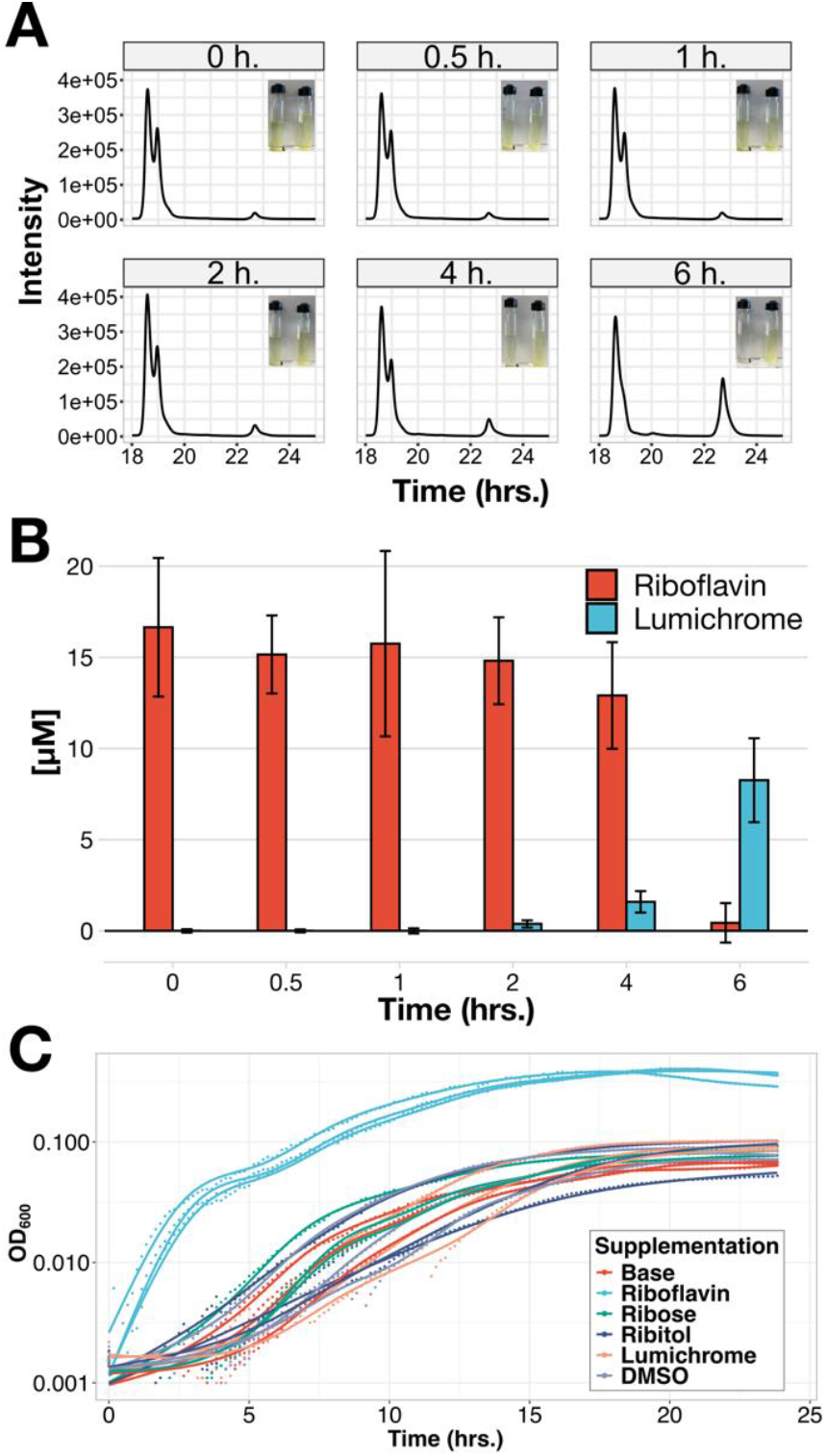
*C. scindens* catabolizes riboflavin to lumichrome, but neither lumichrome nor potential side chain catabolites themselves enhance growth. A) HPLC traces showing riboflavin peak (∼19m) and an unidentified peak (∼23m). Traces are representative of n=3 biological replicates. Representative images of an inoculated culture (left) and control (right) are shown in insets. B) Quantitation of riboflavin and lumichrome in *C. scindens* culture supernatants over time by targeted mass spectrometry. Error bars are 2xSE of n=3 biological replicates. C) Growth curves showing *C. scindens* growth on base medium (0.13µM riboflavin) or medium supplemented with 17.5µM riboflavin (teal), 200µM ribose (green), 200µM ribitol (dark blue), 200µM lumichrome (salmon), or a DMSO control (light blue). Each trace is one biological replicate (n=3); lines are non-parametric smooth fits (generalized additive model).

To determine whether *C. scindens* degraded riboflavin, we performed HPLC on supernatants from Hungate tube cultures sampled at between zero and six hours of growth (see Methods). We identified a peak as riboflavin at 17 minutes, which decreased over this time period; an unknown peak 4 minutes later had the opposite trend, increasing exponentially in height over the course of six hours (Figure 4B). We noted that the unknown peak was also present at a low level at time zero; commercially available riboflavin typically contains low level of breakdown products that form upon exposure to light and oxygen. Depending on the conditions, these products can include lumichrome and lumiflavin.

We therefore tested whether lumichrome, lumiflavin, and/or isoalloxazine were being formed during growth, using liquid chromatography-mass spectrometry with fragmentation (LC-MS/MS). Of these, only lumichrome was detected, and it was detected at the same retention time as our standard in media, indicating that it was not an in-source fragment of another metabolite. When quantitated over the course of *C. scindens* growth, it showed the expected trend based on the HPLC data (Figure 4C), indicating that lumichrome was indeed the major product. Approximately half of the provided riboflavin (in moles) appeared to be catabolized to lumichrome over the first six hours of growth.

The only characterized bacterial pathways for degrading riboflavin to lumichrome are from *Microbacterium maritypicum*^34^ and closely related *Devosia riboflavina*^35^ (formerly *Pseudomonas riboflavina*). However, both require oxygen for riboflavin cleavage, which is catalyzed by the monooxygenase RcaE. The pathway does not proceed anaerobically even when reconstituted *in vitro*. As *C. scindens* is a strict anaerobe, it is unsurprising that a BLASTP search failed to find any significant similarity to RcaE in its genome. This is also consistent with the reported taxonomic distribution of the riboflavin degradation cluster, which appears to be specific to the Actinomycetota and the Alphaproteobacterial genus *Devosia*^35^. There have been reports of gut microbes that degrade riboflavin to lumichrome^57^, but these strains were never identified and appear to have been lost. Interestingly, one anaerobe isolated from goat rumen, *Faecalicatena fissicatena*, also a member of the *Lachnospiraceae*, has been shown to degrade riboflavin, but to a different major product, hydroxyethylflavin (HEF, or “ethanol lumiflavine”)^58^. This is therefore the first report of a taxonomically-identified microbe capable of degrading riboflavin to lumichrome anaerobically.

### Neither lumichrome nor predicted side-chain catabolites enhance *C. scindens* growth

Riboflavin has a ribityl chain attached to its isoalloxazine ring system, which is missing in lumichrome. *M. maritypicum* and *D. riboflavina* cleave this chain using an oxidative process that releases this chain as ribose, which can then serve as a carbon and/or energy source^34,35^. One possibility is therefore that riboflavin enhances *C. scindens* growth by providing an alternative carbon source. While this carbon source would be much lower in concentration than the fructose in our medium (present at 125X the concentration of riboflavin), it is still possible that it could contribute disproportionately to growth by allowing the fructose to be used more efficiently. Another possibility is that lumichrome, rather than riboflavin, could be the molecule that actually enhances *C. scindens* growth, especially as lumichrome is present in trace amounts in commercially-available riboflavin because of oxidative photodegradation.

To assess these possibilities, we grew *C. scindens* in DM with an excess (200µM) of ribose, ribitol (the expected product of non-oxidative cleavage), or lumichrome. Lumichrome is much more hydrophobic than riboflavin, limiting its solubility, so we prepared its stock solution in DMSO and compared it to a vehicle control. We saw no growth enhancement in any condition other than 17.5µM riboflavin supplementation (Figure 4D), even though the *C. scindens* genome does encode predicted homologs of the *rbsABC* ribose import system. This indicates that riboflavin itself enhances *C. scindens* growth, and does so by a mechanism other than providing ribose or ribitol.

### A specific gene neighborhood is induced during *C. scindens* growth on high riboflavin

Since the above experiments pointed to a presently unknown pathway for riboflavin degradation, we performed transcriptomics on *C. scindens* cultures during log-phase growth and stationary phase, in either supplemented (17.5µM) or base (0.13µM) riboflavin. Most transcripts had similar trends that were driven mainly by the phase of growth (Figure 6A). However, a small subset of transcripts did show strong, statistically-significant differential expression across media (Figure 6B, p_adj_≤0.05 and log_2_(fold change)>2, DESeq2 Wald test).

Both predicted riboflavin transporters (*ribU1* and *ribU2*) were upregulated in the base medium compared to the supplemented medium, providing additional evidence that these indeed function in riboflavin import. Additionally, one predicted homolog of the Bacillus stress response gene *ydaJ* was highly correlated with *ribU1/2* expression. While the function of YdaJ is unknown, we noted that it is a predicted flavin-dependent pyridoxamine-5-phosphate (PNP) oxidase, taking PNP to the active pyridoxal-5-phosphate (PLP), and would therefore bind both FAD and pyridoxamine.

The genes upregulated under riboflavin supplementation, in contrast, were almost all located in a separate, single neighborhood in the *C. scindens* genome (Figure 6C). Coverage plots indicated that these likely formed at least three distinct transcriptional units (Supplementary Figure S1). As most of these genes had no annotated name, we have labeled them here as *finA-K*, for “flavin induced neighborhood.” This neighborhood encoded a predicted homolog of the bifunctional riboflavin kinase/FMN adenylyltransferase RibF (*finB*), as well as a predicted Class I aldolase (*finC*), an alcohol dehydrogenase (*finD*), an aldehyde dehydrogenase (*finG*), and two genes homologous to glycine/betaine/sarcosine reductase subunits (*finHI)*.

Because *F. fissicatena* was previously reported to catabolize riboflavin to the related product HEF, we also used MMseqs2 to identify reciprocal best hits between *F. fissicatena* and *C. scindens* coding sequences. The results revealed that *F. fissicatena* indeed encoded a similar neighborhood, but lacking the glycine/betaine/sarcosine reductase homologs. One possible explanation is therefore that HEF production may occur through a variant of the same pathway.

### Orthologous neighborhoods can be detected across related *Lachnospiraceae*

We next performed a genomic analysis to identify other *Lachnospiraceae* with the same gene neighborhood, and to compare its distribution to riboflavin auxotrophy and transport (Figure 5D). Using the OrthoFinder pipeline^59^, we built a tree of 792 high-quality *Lachnospiraceae* isolate and metagenomically-assembled genomes from the Genome Taxonomy Database (GTDB), including the genomes of the five strains we tested in our initial experiment plus one outgroup (*Vallitalea_A okinawensis*); we also assembled orthogroups and constructed orthogroup gene trees.

**Figure 5:**
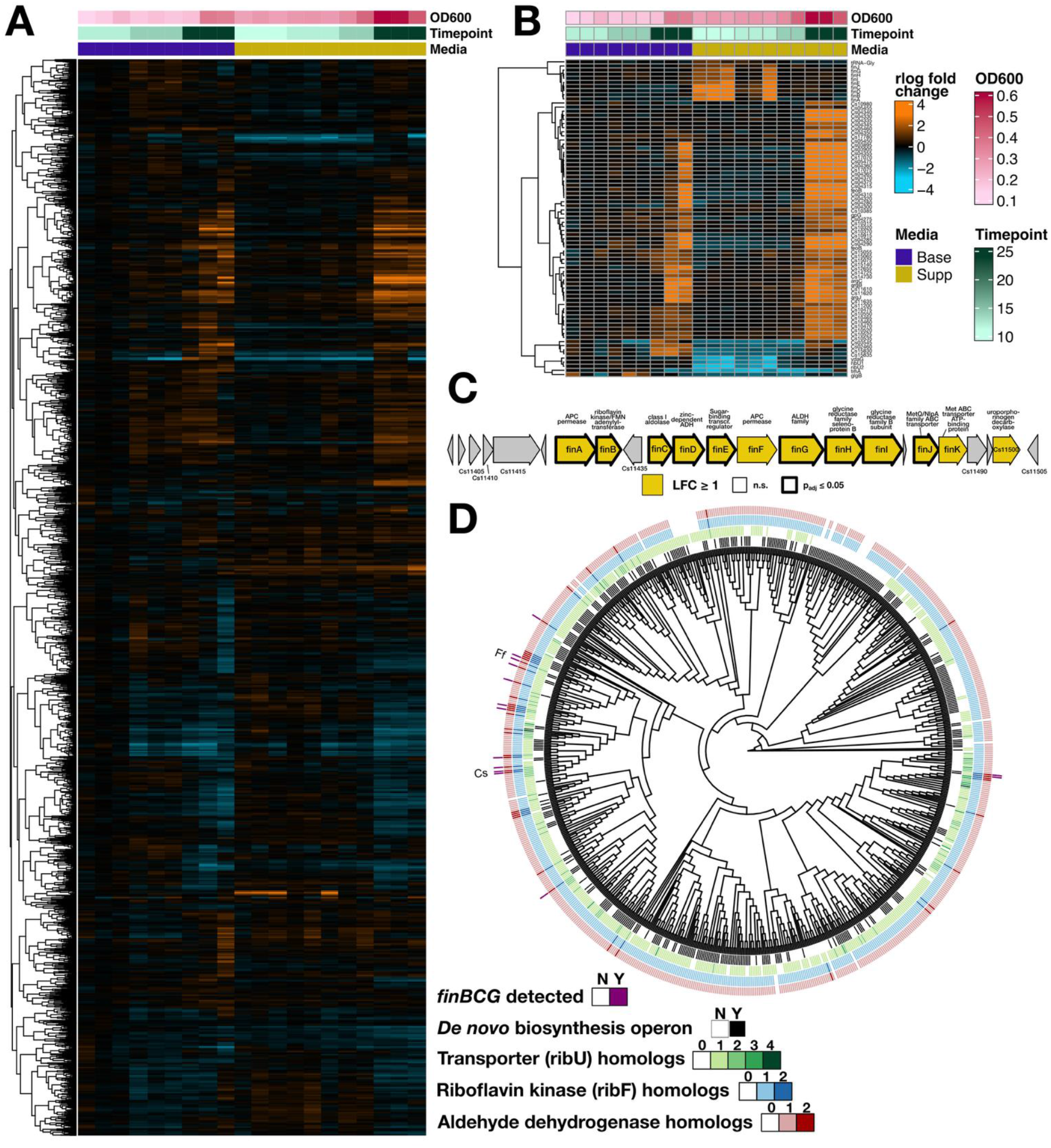
A single gene neighborhood, with sporadic distribution across *Lachnospiraceae*, is induced on high riboflavin. A) Clustered heatmap showing all transcripts under base media (0.13µM riboflavin, blue) and supplemented (17.5µM, orange). Counts were rlog-normalized using DESeq2. B) Transcripts with significant (adjusted p ≤ 0.05, log_2_ fold-change > 2) effects by media. C) *C. scindens* genomic region containing most induced genes. Genes are highlighted in orange if they had a log_2_ fold-change of at least 1, and outlined in black if they were found to be significant (by the criteria in B). D) Tree of *Lachnospiraceae* reference genomes showing, from inside to outside, distribution of de novo riboflavin biosynthesis operon (black), number of *ribU* homologs (olive), number of *ribF* homologs (blue), number of aldehyde dehydrogenase homologs (maroon), and whether or not at least 8 of the unique orthogroups from the region shown in C could be detected (navy). Taxon labels: Cs, *Clostridium scindens*; Ff, *Faecalicatena fissicatena*.

We found that the *de novo* riboflavin biosynthetic pathway had a patchy, heterogeneous distribution among the *Lachnospiraceae*, with 49% of the *Lachnospiraceae* predicted to be auxotrophs based on lacking this pathway (Figure 5D, black lines). Not all *Lachnospiraceae* encoded *ribU*, but matching a previous observation^16^, the *ribU* transporter gene was almost always present when the *de novo* pathway was missing. Furthermore, as we saw in *C. scindens* and *B. coccoides*, many other *Lachnospiraceae* had a second *ribU* paralog (12.5%), with some diverged *Lachnospiraceae* having even a third or fourth (Figure 5D, amber colors).

Several of the genes in the *fin* neighborhood were assigned to very large orthogroups that contained multiple paralogs per genome, making it difficult to assess which genomes truly had orthologous neighborhoods. For example, most alcohol dehydrogenases were grouped into a single large protein family, likely representing proteins with many different specific substrates. We therefore used the gene trees to refine three of these larger protein families (riboflavin kinase/FMN adenylyltransferase, class I aldolase, and aldehyde dehydrogenase) into sets of specific orthologs, using the *C. scindens* and *F. fissicatena* sequences as guides.

Notably, in all three cases, the *C. scindens* and *F. fissicatena* sequences formed part of a monophyletic clade that diverged sharply from other sequences in the orthogroup (Supplementary Figure S2: Orthogroup gene trees for three genes in the fin neighborhood: A) bifunctional riboflavin kinase/FAD synthetase (*finB*), B) aldolase (*finC*), and C) aldehyde dehydrogenase (*finG*). The *C. scindens* and *F. fissicatena* orthologs are labeled, and the clades selected for identifying potential orthologous neighborhoods are highlighted in yellow.). This clear separation, consistent with functional divergence, allowed us to use membership in these subclades to call orthologs. We provisionally classified genomes that had orthologs of all three of these proteins as *fin+* (Supplementary Table S5: Table of all potentially orthologous *fin* neighborhoods detected.).

This analysis yielded 12 other representative genomes besides *C. scindens* and *F. fissicatena* that encoded *fin* orthologs, including *B. coccoides* (Figure 5D, navy lines). The *fin* orthologs had a sporadic distribution across *Lachnospiraceae*, possibly indicating horizontal transfer, but were most concentrated in a subclade that included the genera *Dorea, Mediterraneibacter*, and *Muricomes*. Some genomes had a more *C. scindens*-like neighborhood, containing genes in the glycine reductase family (e.g., *Muricomes intestini*), while others had a more *F. fissicatena-*like neighborhood lacking these (e.g., *Mediterraneibacter hominis*). The *F. fissicatena* and *M. hominis* neighborhoods were both flanked by another aldehyde-alcohol dehydrogenase gene that was not in the same orthogroup as any of the original *fin* genes. Thus, at least two variants appear to be carried by multiple *Lachnospiraceae* (Supplementary Figure S3).

### Genomic and metabolomic data point to gut Gram-negatives as a potential riboflavin source for *Lachnospiraceae in vivo*

If *C. scindens*, and potentially other *Lachnospiraceae*, are dependent on a vitamin they also degrade, then we would expect that vitamin to be abundant in its local context. However, in mammals, dietary riboflavin is known to be efficiently absorbed in the upper small intestine. Some have argued that unabsorbed riboflavin could persist in the large intestine^60^, but according to recent survey data^61^, the average American over twenty years old consumes only 1.92 ± 0.034 mg of riboflavin daily, well below the estimated maximum of 27 mg that could be absorbed in a single dose^62^. This suggests that either *C. scindens* is metabolizing riboflavin that crosses from the blood to the colon (as is observed for other bloodborne nutrients like lactate and urea^10^), or there is an alternative source of riboflavin in the gut, possibly microbial in origin.

We re-analyzed metabolomic data from a study^63^ that compared germ-free to conventionally colonized mice. Riboflavin was found to be much more abundant in colonized mouse intestine, cecum, and/or feces compared to germ-free mice (Figure 6A). This is consistent with gut microbes overproducing riboflavin *in vivo*. Notably, Han et al.^63^ also sampled serum, where a modest but significant trend in the opposite direction was observed: serum riboflavin was highest in the germ-free mice and lower in colonized mice (Figure 6A). This suggests that microbial riboflavin is not substantially absorbed by the host. Finally, Han et al. also measured a metabolite that they identified as lumichrome, and this metabolite was highly associated with colonized mouse cecum/feces (Figure 6A). Since riboflavin can be degraded to lumichrome in the presence of UV light and oxygen, we cannot exclude the possibility that the reported lumichrome concentrations may mainly reflect riboflavin abundance; however, this is consistent with previous results in rabbits, where HEF was found only in colonized and not germ-free animals^38^.

**Figure 6:**
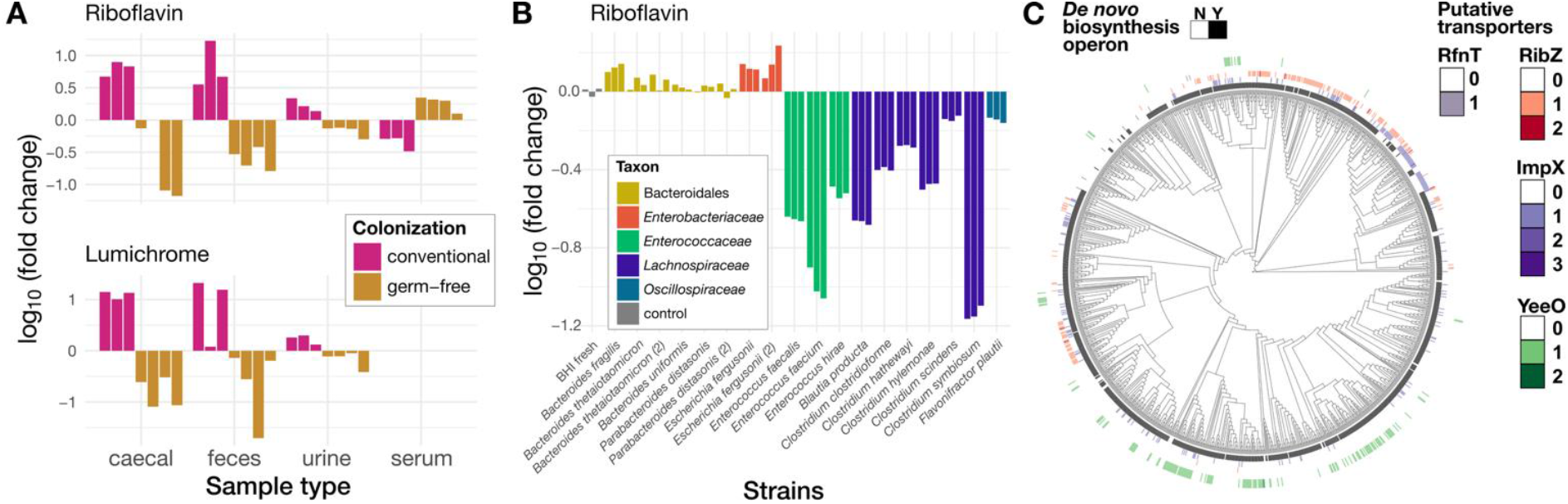
Metabolomic and genomic data support a role for riboflavin cross-feeding in the gut. A) Metabolomics data from Han et al.^63^ showing in vivo measurements of riboflavin (top) and lumichrome (bottom) across four different tissues in colonized (navy) and germ-free (orange) mice. Measurements were normalized to the average of each group to show log-fold changes. B) Metabolomics data from Ho et al.^64^ showing riboflavin concentrations in spent media (BHI) for gut isolates from several different families (colors). Uninoculated BHI is shown in grey and was used to normalize the other measurements. C) Tree of Bacteroidales genomes with the riboflavin *de novo* pathway (black) and copies of putative transporters (RfnT homologs, teal; RibZ, red; ImpX: blue; YeeO, green) highlighted in rings; note that YeeO is a potential exporter.

To determine which microbes could be the source of this riboflavin, we turned to metabolomic data from a recent set of *in vitro* culture experiments that sampled several diverse gut isolates grown in rich media (BHI), including members of the genera *Escherichia* and *Bacteroides*^64^. Riboflavin in spent media from *B. fragilis, E. fergusonii*, and to some extent *B. thetaiotaomicron* was indeed higher after culturing than in uninoculated controls (Figure 6B). Further supporting the idea that many *Lachnospiraceae* are riboflavin auxotrophs and/or have a high requirement for this vitamin, all *Lachnospiraceae* tested in this study depleted significant quantities of riboflavin from BHI, while none of the tested Gram-negatives did so.

We then compared these patterns to those obtained from an analysis of 1,291 genomes (Supplementary Table S6) from the most abundant Gram-negative clade, the *Bacteroidales*, plus three outgroups (*Flavobacterium psychrophilum, Flavobacterium columnare*, and *Flavobacterium johnsoniae*). In contrast to the *Lachnospiraceae*, we found that *de novo* biosynthesis was highly conserved in this clade, and riboflavin import was much less so. However, some predicted transporters had a sporadic distribution, suggesting transfer into the clade (Figure 6C). Interestingly, *Bacteroidaceae* specifically were more likely to have homologs of the *E. coli* MATE transporter YeeO, which has been shown in *E. coli* to be a flavin exporter^65^. Further work will be required to confirm the function of this homolog in *Bacteroidaceae*.

Overall, these observations are consistent with a model where *Bacteroidaceae* and *Enterobacteriaceae* overproduce riboflavin in the gut, which is then taken up by *Lachnospiraceae*, and in some cases, is finally degraded to either HEF or lumichrome.

## Discussion

### Anaerobic catabolism of riboflavin by *Lachnospiraceae*

We found that *C. scindens* growth was enhanced by riboflavin supplementation, that this enhancement continued up to at least 17.5µM, and that *C. scindens* also degraded approximately half of the riboflavin in the medium to lumichrome over the course of six hours of log-phase growth. Furthermore, a single gene neighborhood was strongly induced on high (17.5µM) riboflavin, relative to a 0.13µM media control. While it has previously been suggested that lumichrome is a gut bacterial metabolite and not a host-derived molecule^66^, this is the first report to identify a specific microbe, *C. scindens*, that is capable of degrading riboflavin to lumichrome without the use of oxygen.

To our knowledge, the only other specific anaerobe that has been reported to degrade riboflavin is an isolate of *Faecalicatena fissicatena* collected circa 1970, which produced the alternative product hydroxyethylflavin (also HEF, or “ethanol lumiflavine”)^67^. This isolate of *F. fissicatena* also produced color changes on high-riboflavin media, though these were different from what we observed (green and crimson, changing to orange on oxygen exposure, as opposed to the color dissipation). Interestingly, we found that a shorter variant of the *fin* neighborhood, lacking two genes in the glycine reductase family, was also present in *F. fissicatena*. The existence of these pathway variants could also potentially explain why previous studies have found multiple related breakdown products of riboflavin, including HEF and formylmethylflavin (FMF), in, e.g., human urine^36,37^ or the ceca of colonized rabbits^38^.

We did observe homologs of the *fin* genes in *B. producta*, and like *C. scindens*, it appears to be a riboflavin auxotroph based on its genome content. However, riboflavin supplementation did not restore its growth. Future work is required to determine the growth factors it may be missing on our defined medium, or alternatively, inhibitory factors that may be present.

### Why does *C. scindens*, a riboflavin auxotroph, degrade riboflavin?

Ribose, ribitol, and lumichrome failed to enhance *C. scindens* growth. Therefore, while *C. scindens* may still metabolize the sugar (or sugar alcohol) that is cleaved from riboflavin during lumichrome formation, the utilization of these sugars does not appear to explain the large growth benefit we observed.

One possible explanation could be a type of directed overflow metabolism^68^ for reduced flavins, potentially carried out by the products of the flavin-induced neighborhood (*fin*) that we identified here. During carbon oxidation, FAD and FMN are reduced to FADH_2_ and FMNH_2_; in anaerobes, these are typically re-oxidized via pathways that include donating electrons to hydrogenase (carried by flavodoxins), or to substrates like fumarate^69^, urocanate^70^, or, in the case of *C. scindens*, bile acids^71^. In the absence of a suitable electron acceptor, oxidized flavins may be unavailable for key central carbon metabolic enzymes such as, e.g., pyruvate-flavodoxin/ferredoxin oxidoreductase (PFOR). An excess of reducing equivalents, or “reductive stress,” could also have other negative downstream effects^72^. Flavin biosynthetic intermediates are also known to be targeted for overflow metabolism because of their reactivity^32,73^. In the presence of abundant oxidized riboflavin, therefore, it could be advantageous to simply catabolize the excess reduced flavins into a product like HEF or lumichrome (for which the concentration gradient would likely favor non-energetic export), then import “fresh” oxidized riboflavin for FMN and FAD synthesis.

The first step of a flavin overflow pathway would likely start with catabolizing FADH_2_ or FMNH_2_. We noted that the *fin* neighborhood contains a bifunctional FAD synthetase. While this enzyme is named for its ability to convert riboflavin to FMN and then to FAD, some prokaryotic bifunctional FAD synthetases also have FAD pyrophosphorylase activity, meaning that they can also make FMN from FAD. Furthermore, one such bifunctional FAD synthetase, purified from *Streptococcus pneumoniae*, was shown to be highly selective for de-adenylylating the reduced form, FADH_2_ ^74^; a similar preference has also been observed in a *B. subtilis* FAD synthetase^75^. Notably, this reverse reaction can also generate ATP from pyrophosphate. An interesting future direction would therefore be to test whether FinB indeed functions as an FADH_2_-specific pyrophosphorylase.

Another observation supporting the idea of flavin overflow has to do with the lumichrome produced by *C. scindens*. Oxidized lumichrome, like riboflavin, is bright yellow, and because of its lower solubility, aerobic catabolism of riboflavin to lumichrome typically yields a bright yellow precipitate, as observed for *M. maritypicum*^34^. However, we did not observe this in our *C. scindens* cultures. Fully reduced lumichrome, in contrast, absorbs mostly in the UV spectrum^76^ and therefore has little visible color, meaning that the lumichrome being formed in our culture supernatants could be in the reduced form.

An alternative possibility is that the anaerobic riboflavin degradation to lumichrome that we observe could be due to metabolic damage^77^, with the *fin* neighborhood playing a separate role in flavin metabolism. Cofactors can be damaged by either non-enzymatic spontaneous or enzymatic side reactions^78^. Unlike, e.g., nicotinate cofactors, flavins can exist in either singly- or double-reduced forms, making reactions that depend on radical chemistry more likely. Certain flavin biosynthesis intermediates are indeed known to be labile^32,73^, and in the most famous example of non-enzymatic degradation, riboflavin can decompose in the presence of light and oxygen. Photolysis cannot explain our current results (note that we did not observe bleaching of uninoculated controls), but does speak to the general reactivity of flavins.

On the other hand, despite this potential for reactivity, pulse-chase labeling experiments in prototrophic *E. coli* and *B. subtilis* strains have found low baseline rates of flavin damage. In these experiments, the “pool turnover” rate of cofactors was measured, and FAD and FMN were actually found to be some of the most stable (compared to, e.g., folates or ATP), with most of the turnover accounted for by dilution caused by cell division.^79^ What difference in the cellular environment and/or growth conditions of *C. scindens*, then, would favor much higher rates of flavin damage, with nearly half the riboflavin converted into lumichrome? One possibility is that the above pulse-chase experiments were performed aerobically, and therefore also in the presence of a good electron acceptor. It remains to be determined whether flavin pool turnover in a prototroph would remain this low under anaerobic and/or reductive stress conditions, which might accelerate metabolic damage.

### Vitamin catabolism could be an underappreciated contributor to the gut microbial food web

Our reanalysis of previous metabolomic data shows that riboflavin is likely produced by gut microbes, and also suggests that it may be cross-fed from Gram-negative to Gram-positive bacteria. Most previous studies of vitamin metabolism in gut microbes have focused on auxotrophy as a main potential driver of cross-feeding interactions. Vitamin catabolism, in contrast is largely understudied; as mentioned above, the aerobic pathway for riboflavin degradation was only elucidated more than 70 years after it was first described in soil isolates. Catabolism could potentially be relevant beyond riboflavin metabolism, as other vitamins like pyridoxal, nicotinate, and especially biotin are also elevated in colonized vs. germ-free mouse guts^80^. Other B vitamins with *in vitro* evidence for gut microbial production and uptake include thiamine^18,81^ and folate^18^.

While our supplementation experiments were performed with 17.5µM riboflavin, the concentration *in vivo* may not need to be as high if flavins are being continuously secreted by other microbes. Notably, microbes such as *E. coli, P. fluorescens*, and *B. subtilis* have all been observed to produce more flavins than they need, and to excrete flavins into the supernatant^50^. One reason may be that flavin production is not tightly regulated at the level of feedback inhibition^50^, perhaps because flavin biosynthetic intermediates can be toxic to the cell^73^. Alternatively or in addition, *Lachnospiraceae* could actively elicit overproduction in other microbes. While this could be an active process (e.g., via signaling), it could also simply be that lowering the extracellular concentration of riboflavin in the local environment (and replacing it with lumichrome) could favor export by producers.

### Riboflavin degradation products may have further impacts on both microbiome and host

There is precedent for microbial sensing of lumichrome: for example, the LasR quorum sensing receptor from P. aeruginosa, whose preferred ligand is an acyl-homoserine lactone, can also be activated by both riboflavin and, especially, lumichrome, though this activation is weaker than for the preferred ligand^82^. *P. aeruginosa* is also found in soils; production of lumichrome is known to be an important function of rhizosphere bacteria (potentially including species of *Devosia*, as some of these aerobically catabolize riboflavin to lumichrome), as lumichrome stimulates plant root growth^83^. Finally, a pathway for the complete degradation of lumichrome to organic acids and ammonia was just identified in *Nocardioides simplex*, although this pathway is also dependent on oxygen^84^; it is therefore possible that lumichrome is not only sensed, but further metabolized in the gut.

Perhaps even more importantly, in addition to its role as a vitamin and cofactor component, riboflavin and related metabolites are known to be sensed by the mammalian immune system. Mucosal-associated invariant T cells, which are enriched in the mammalian gut, express a specific narrow range of T-cell receptors (TCRs) that bind a particular MHC class I-like antigen-presenting protein, MR1. MR1 has been shown to bind small molecules related to B vitamins^85^, and in particular, when bound to adducts of riboflavin biosynthetic intermediates, MR1 activates MAIT cells^86^. Since humans do not synthesize riboflavin *de novo*, these intermediates are necessarily microbial in origin. Furthermore, MR1 bound to riboflavin itself inhibits MAIT cell activation; as colonized guts are much higher in riboflavin, this may help to control MAIT cell activation.

How do flavin catabolites fit into this picture? Interestingly, a recent study showed that lumichrome, as well as other breakdown products such as FMF and lumiflavin, actually bind MR1 with higher affinity than riboflavin, but unlike the biosynthetic intermediates, also inhibit MAIT cell activation^87^. The study authors explained this phenomenon by referring to lumichrome as a “host metabolite,” pointing out that it is found in human blood. However, studies on rat tissue have argued against an animal origin of lumichrome^66^, and as we have demonstrated, human gut microbes can catabolize riboflavin to lumichrome. Others have implicated gut microbes in the degradation of riboflavin to products like HEF^38^ and FMF^36^, noting that (as we found in our re-analysis of metabolomic data) riboflavin catabolites are enriched specifically in colonized vs. germ-free animals. An alternative interpretation of the above findings is therefore that commensal bacteria that catabolize riboflavin inhibit MAIT cell activation, which could be another non-SCFA-dependent mechanism by which they contribute to an anti-inflammatory tone in the gut. Future work, potentially in a gnotobiotic system, would be required to directly test the *in vivo* impact of riboflavin degradation.

## Materials and Methods

### Bacterial strains

Initial growth curve experiments were performed with five representative members of the *Lachnospiraceae* family: *Robinsoniella peoriensis* B-23985, *Anaerostipes caccae* DSM 14662, *Blautia coccoides* ATCC 27340, *Dorea formicigenerans* ATCC 27755, and *Clostridium scindens* ATCC 35704. Strain designations, culture collection sources, and accession numbers are provided in Supplementary Table S1. All strains were verified by amplifying the 16S rRNA gene using bacterial universal primers 27F and 1492R, followed by Sanger sequencing, and maintained as glycerol stocks (16.6% v/v) at −80°C.

### Pre-culture conditions

Strains were struck onto agar plates containing brain heart infusion supplemented with hemin, menadione, cysteine, and resazurin (BHI-CHK), and incubated anaerobically inside a Coy anaerobic chamber maintained with an atmosphere of 85% nitrogen, 10% carbon dioxide, and 5% hydrogen. Because hydrogen is consumed by the catalyst, the working hydrogen concentration was typically 2-3%. Three independent colonies per strain were inoculated into BHI-CHK broth and grown anaerobically for 24h prior to transfer into a chemically defined medium.

### Chemically defined medium formulation

A chemically defined medium (DM) was adapted from Devendran et al.^41^ with some modifications. We chose fructose as the carbon source because preliminary analysis of RefSeq and KEGG annotations showed that fructokinase and fructose-specific PTS transporters appeared to be common among *Lachnospiraceae*, and also because fructose is a common component of dietary fibers like inulin. We also used 100X vitamin and mineral stock solutions, which were based on Wolin and Wolfe’s formulation^88^. For minerals, we used a commercial supplement (ATCC, MD-TMS) that used EDTA as a chelator instead of nitrilotriacetic acid. We prepared our own vitamin stock solutions; differing from Wolin and Wolfe’s original formulation, we provided vitamin B_6_ in three forms, pyridoxin, pyridoxal and pyridoxamine, as some *Lachnospiraceae* lacked the predicted enzymes to activate one or more of these to pyridoxal-5-phosphate (PLP). Finally, our medium did not include resazurin.

The final medium therefore consisted of (final concentrations): fructose [25.00 mM]; NaHCO_3_ [45.00 mM]; NaPO_4_ [36.60 mM]; KPO_4_ [22.00 mM]; NaCl [8.556 mM], NH_4_Cl [18.7 mM]; biotin [0.00008 mM]; folic acid [0.00005 mM]; pyridoxin [0.00049 mM]; thiamine [0.00015mM]; riboflavin [0.00013 mM]; nicotinic acid [0.0004 mM]; pantothenic acid [0.00021 mM]; cobalamin [7.40e^-7^ mM]; p-aminobenzoic acid [0.0036 mM]; (DL)-alpha lipoic acid [0.00024 mM]; pyridoxal [0.001 mM]; EDTA [6.8 µM]; MgSO_4_ [48 µM]; MnSO_4_ [12 µM]; FeSO_4_ [1.44 µM]; Co(NO_3_)_2_ [1.36 µM]; CaCl_2_ [3.6 µM]; ZnSO_4_ [3 µM]; CuSO_4_ [0.16 µM]; AlK(SO_4_)_2_ [0.156 µM]; H_2_BO_3_ [0.64 µM]; Na_2_MoO_4_ [0.164 µM]; Na_2_SeO_3_ [0.0232 µM]; NaWO_4_ [0.12 µM]; NiCl_2_ [0.336 µM]; all 20 amino acids [0.2 mM]; Na_2_S [1.2 mM]; and L-cysteine [1.5 mM]. We also prepared a version of this defined medium supplemented with acetate (DM+Ace) by adding sodium acetate at 45 mM.

For experiments conducted inside the anaerobic chamber, all media were prepared aerobically, filter sterilized, and pre-reduced inside the anaerobic chamber at least overnight prior to inoculation.

### Chemically defined media growth curves

Growth assays were conducted in sterile 96-well plates with a final working volume of 200 µL per well. Following pre-culture growth, 2 mL of each BHI-CHK culture was harvested by centrifugation at 4,000 rpm for 10 min inside the anaerobic chamber. Supernatants were decanted, and cell pellets were resuspended in 10 mL of pre-reduced DM. Cultures were inoculated by transferring 20 µL of the resuspended cells into 180 µL of fresh DM. Plates were sealed with a gas-permeable Breathe-Easy film (Diversified Biotech BEM-1) and transferred to an automated plate stacker connected to an Agilent microplate reader. Optical density at 600 nm (OD_600_) was measured every 10 min. for 24 hours. Prior to each measurement, plates were shaken for 30s using a double-orbital motion to homogenize the culture. As it was difficult to perform subcultures from a plate sealed with a Breathe-Easy film without aerosol contamination, a parallel replicate plate was sealed with a Breathe-Easier film, and was incubated adjacent to the plate stacker under the same conditions.

Following 24h of incubation, 20 µL from the replicate plate was transferred into 180 µL of fresh pre-reduced DM into two new 96-well plates. This subculturing procedure was repeated for a total of three sequential transfers under identical anaerobic and measurement conditions. Growth curve parameters (e.g., carrying capacity, growth rate) were fit to a logistic equation using GrowthCurver^89^.

### Comparative genomics in cultivated strains

Genomes from *D. formicigenerans, C. scindens, B. coccoides*, and *A. caccae* were obtained from the NCBI Refseq database. In addition, the genome from *R. peoriensis* was sequenced and assembled as part of this study, using a combination of short-read Illumina and long-read Oxford Nanopore sequencing.

Assembly was performed according to the guidelines of Wick, Judd, and Holt^90^. Raw short-reads for *R. peoriensis* were processed using fastp^91^ v0.24. The raw long-reads were subsampled into independent read sets with Trycycler^92^ v0.5.5 then assembled with Flye^93^ v2.9.3, Minipolish v0.1.3^94^, and Raven^95^ v1.8.3. Each assembly was clustered and reconciled with Trycycler and contigs were manually reviewed. Six contigs were excluded due to large gaps and poor alignments, indicating large structural errors. All contigs that passed a manual check were then processed with multiple sequence alignment and a consensus was generated with Trycycler. The final consensus was polished with Medaka^96^ v2.0.0 using long-reads and further refined with short-reads using Polypolish^97^ v0.5.0. Finally, one additional round of polishing was completed with Polca^98^ v0.2.1.

All genomes were annotated using the anvi’o platform v8^99^. KEGG Orthologs (KOs) were assigned using the anvi-run-kegg-kofam workflow, which identifies KOs with a Hidden Markov Model approach, using adaptive cutoff adjustment to improve sensitivity (KEGG release 107.0 September 2023).^46^ The default parameters were used unless otherwise specified.

A binary presence-absence matrix of KOs was generated from anvi’o annotations for all genomes, with KOs scored as present if at least one copy was detected in a given genome. The resulting matrix was used for comparative genomic analyses across strains.

### Riboflavin supplementation

To assess the effect of the riboflavin availability on growth, the chemically defined medium was supplemented with riboflavin at 17.5 µM. We used a saturated stock solution of riboflavin in water, heating to 60ºC for 4-6m before filter sterilizing. Using absorbance to measure the concentration, we found that 175 µM riboflavin remained in solution after sterilization, consistent with the reported solubility limit in water. This stock solution was made fresh on the day of each experiment in order to avoid possible degradation. We used this stock at 10X to yield media with 17.5 µM riboflavin. Growth curve experiments were performed inside the Coy anaerobic chamber; growth curve experiments, incubation, and data acquisition were performed using the same protocol as described in previous sections.

### Growth with riboflavin degradation products

To evaluate whether potential degradation products of riboflavin could support growth, we supplemented the chemically defined medium with ribose, ribitol, or lumichrome at a final concentration of 200 µM. Because of its low solubility in water, lumichrome stock solutions were prepared in DMSO, so we additionally performed a DMSO vehicle control. Growth curves were performed under strict anaerobic conditions inside the Coy anaerobic chamber using a microplate protocol, inoculation strategy, and incubation conditions as described above. Optical density at 600 nm (OD600) was measured using an Alto plate reader (Cerillo) without shaking.

### Growth in Balch/Hungate tubes and measurement of hydrogen production

To evaluate the growth under 100% CO_2_, growth experiments were conducted using Balch and Hungate tubes outside the anaerobic chamber. Chemically defined media was prepared and described as above, either with 17.5µM or 0.13µM of riboflavin. After preparing the media, 9.5 mL were dispensed into Hungate or Balch tubes. All tubes were sealed and sparged with 100% CO_2_ for 20 minutes to establish anaerobic conditions and remove residual oxygen. Following gas exchange, tubes were stored at 4ºC until use.

*Clostridium scindens* was grown under 100% CO_2_. Three independent colonies were picked from BHI-CHK plates and inoculated into BHI-CHK broth inside the anaerobic chamber, followed by incubation at 37ºC for 24 h. After incubation, 2 mL of culture were harvested by centrifugation at 4,000 rpm for 10 min inside the anaerobic chamber. Supernatants were decanted, and cell pellets were resuspended in 10 mL of pre-reduced chemically defined media and transferred into sterile Hungate tubes prior to removal from the chamber.

For inoculation, 0.5 mL of the resuspended culture were transferred into prepared Hungate and Balch tubes using a sterile 1 mL syringed fitted with 23G needle. Following inoculation, tubes were incubated statically at 37ºC in the dark. Growth was monitored by measuring OD600 at hourly intervals through the tubes.

Following completion of growth measurements, hydrogen production was quantified from the headspace of the Hungate and Balch tubes using gas chromatography. Headspace samples (250 µL) were collected from each tube using 1 mL gas-tight syringe and injected into a gas chromatography equipped with a thermal conductivity detector (TCD) and helium as the carrier gas. Hydrogen separation was achieved using a molecular sieve column. Uninoculated medium controls were processes in parallel to account for background hydrogen levels.

### Extracellular sampling and analysis

To quantify and characterize metabolites over time, *C. scindens* was grown in Hungate tubes under anaerobic conditions outside of the Coy anaerobic chamber, using the same media preparation and inoculation strategy, and 100% CO_2_ as described above. Chemically defined media was supplemented with 17.5µM of riboflavin.

Cultures were inoculated by transferring 500µL of resuspended cells into Hungate tubes. At different time points following inoculation (0, 0.5, 1, 2, 4, and 6 hrs), 1.5 mL of cultures were collected from individual tubes. Samples were then centrifuged at 15,000 rpm for 5 min, and the supernatant was collected and immediately snap frozen for downstream ultra-fast liquid chromatography (UFLC) analysis. Following sample collection, tubes were discarded and not reused, resulting in destructive sampling each time point. All experiments were performed using three biological replicates. Tubes and supernatants were maintained in the dark, with care taken to minimize light exposure during sampling and measurement, in order to avoid photodegradation of flavins.

Frozen supernatant samples were thawed on ice prior to analysis. For each sample, 200µL of supernatant were mixed with 300µL of ultra-pure water to reduce samples viscosity, then transferred into LC vials.

HPLC analysis was performed on a Shimadzu Prominence system with UV detection at 260nm and 215nm wavelengths. Compounds were resolved using a MacMod Altima C18 reverse phase column at 30ºC. The mobile phase consisted of water containing 0.025% trifluoroacetic acid (Buffer A) and acetonitrile containing 0.025% trifluoroacetic acid (Buffer B). Separation was carried out at a flow rate of 1mg/min from 0%B to 45%B over 15 minutes after an initial hold at 0%B for 5 minutes. A total volume of 480µL was injected for each HPLC run.

Riboflavin and lumichrome were quantified by liquid chromatography–mass spectrometry (LC–MS/MS) using a Sciex Triple Quad 3500 mass spectrometer coupled to a Shimadzu HPLC system. Separation was achieved on a reversed-phase C18 column (Synergi Fusion-RP, 80 Å, 4 µm, 50 × 2.0 mm; Phenomenex) maintained at 40°C, with a flow rate of 0.2 mL min^−1^ and sample injection volume of 10 µL. The mobile phase consisted of water containing 0.025% trifluoroacetic acid (solvent A) and acetonitrile containing 0.025% trifluoroacetic acid (solvent B). The gradient program was 0 min, 100% A; 1 min, 5% B; 9 min, 50% B; 10–11 min, 75% B; and 12–14 min, 5% B. Detection was performed using positive electrospray ionization with multiple reaction monitoring. Riboflavin was monitored at m/z 377→243 (declustering potential 65 V; collision energy 32 V) and eluted at 6.3 min, while lumichrome was monitored at m/z 243.2→198 (declustering potential 124 V; collision energy 30 V) and eluted at 7.6 min.

### RNA sample collection, extraction and sequencing

RNA sequencing experiments were conducted using *C. scindens* grown under the same anaerobic conditions and experimental setup described above for the tube-based growth experiments under 100% CO_2_. Cultures were prepared, inoculated, and incubated identically, except that the experiments were performed exclusively in Balch tubes. Cultures were incubated statically at 37ºC. Sampling time points were selected to capture transcriptional dynamics across growth phases. Samples were collected at 11, 12, 14, and 24 h. following inoculation. At each time point, 5 mL of culture were collected using a sterile syringe fitted with 18G needle and immediately transferred into 10 mL of RNA protect reagent to stabilize RNA. Samples were mixed thoroughly and processed according to the manufacturer’s instruction. Total RNA was extracted using the RNeasy Mini Kit following the manufacturer’s protocol. The base media samples at 11 h post-inoculation had very low RNA yield and were dropped going forward. Extracted RNA was stored at −80ºC until downstream analysis.

RNA was quantified by Qubit™ RNA Assay Kit (Invitrogen, Thermo Fisher, Carlsbad, CA, USA), and 425 ng was used for the RapidOut DNA Removal protocol (Thermo Fisher Scientific, Waltham, MA, USA). Genomic DNA was removed by treating samples with 1 µl of RNase-free DNase I and 2 µl 10X DNase Buffer. Incubation and DNase removal were performed according to the manufacturer’s instructions. Purified RNA was used for RNA sequencing as follows. Ribosomal RNA was depleted from the extracted RNA using the Hybridized Probe, rRNA Depletion, and Probe Removal reaction mixtures from the Illumina Stranded Total RNA Prep (Illumina Inc., San Diego, CA, USA). rRNA-depleted samples were purified using a 2.0x sbeadex SAB bead cleanup (Biosearch Technologies, Hoddesdon, UK) with a single 80% ethanol wash. Following cDNA library construction and indexing, the cDNA libraries were quantified using the Qubit™ 1X dsDNA High Sensitivity (HS) Assay Kit (Invitrogen, Thermo Fisher, Carlsbad, CA, USA) and the fragment sizes were assessed using a 4150 TapeStation System (Agilent Technologies, Santa Clara, CA, USA). Samples were then pooled in equimolar amounts and sequenced on an Illumina NextSeq 2000 with paired end 2−150 bp reads at 25 million reads for each sample.

Raw sequencing reads were processed using Trimmomatic^100^ v0.38. Adapter sequences and low-quality bases were removed using the following parameters: LEADING:3, TRAILING:3, SLIDINGWINDOW:4:15, and MINLEN:50. Reads were trimmed based on when the average quality of the bases dropped below a Phred score of 15 and reads lower than 50 bases after trimming were discarded.

Trimmed reads were aligned with the *Clostridium scindens* reference genome^41^ (NCBI RefSeq Assembly: GCF_004295125.1) using Bowtie2 v2.4.1. Alignment output files were written into SAM format. Alignments were sorted based on chromosomal coordinates using Samtools v1.17. Duplicate reads were identified and marked using the MarkDuplicates function in picard v3.0.0, with the alignment sort order set to “coordinate,” then retained for downstream analysis.

Gene-level read counts were aligned to an annotated reference genome using the featureCounts tool from the subread v2.0.8 package. Reads were aligned to genomic features type gene using gene identification as meta-features. A count table was exported as a text file for downstream analysis.

Differential gene expression analysis was conducted using the R package DESeq2^101^ v1.46.0. One sample with fewer than 1e6 reads was discarded, and genes with zero variance were also dropped. For significance testing, timepoints were binned into either early (<18h) or late (>18h). A model with a two-way interaction, with one factor representing early vs. late timepoints and the other representing supplemented vs. base media, was then fit to the count data; genes with either a significant interaction or a significant media main effect (p_adj_ ≤ 0.05, log_2_ fold change ≥ 2, Wald test) were retained. Heatmaps were generated with ComplexHeatmap^102^ v2.24.1; count data were first transformed using the regularized log (rlog) transformation in DESeq2. For visualization, each gene was normalized by subtracting out the mean value at 12 hours in base (0.13µM) medium, so that plotted rlog values would reflect changes relative to this timepoint.

### Comparative genomics across *Lachnospiraceae* and *Bacteroidales*

To expand comparative analyses across the *Lachnospiraceae* family, representative genomes were selected based on genome quality metrics. Only genomes with an estimated completeness greater than 95% and contamination less than 5% were retained for downstream analyses. High-quality genomes meeting these criteria were obtained by searching the Genome Taxonomy Database (GTDB v202.0). We used representative genomes to minimize redundancy and ensure phylogenetic coverage across the family.

Annotation with KOfams and generation of presence-absence matrices was performed as in the previous section. In parallel, genomes were annotated using Prokka^103^ (with gene calling by Prodigal^104^) to generate predicted protein sequences. The resulting amino acid FASTA files were used as input for orthogroup identification using OrthoFinder v2.5.5^59^ to assess gene family conservation and copy number variants across *Lachnospiraceae* genomes. OrthoFinder was also used to infer a species tree phylogeny via FastTree^105^ based on a concatenated MAFFT^106^ alignment of single-copy orthologs, and to produce gene trees of orthogroups via DendroBLAST^107^.

Specific orthologs of *fin* genes in *Faecalicatena fissicatena* and *Muricomes intestini* were identified using MMseqs2^108^ v18.8cc5c to conduct a reciprocal-best-hit (“rbh”) search against *C. scindens* with default parameters. To find other orthologous neighborhoods, we used the trees of the orthogroups containing the bifunctional riboflavin kinase/FAD synthetase (*finB*), class I aldolase (*finC*), and aldehyde dehydrogenase (*finG*). The *C. scindens* sequences and the *F. fissicatena* orthologs identified using MMseqs2 were highlighted in the tree and used as a guide to divide the tree (see Supplementary Figure S2).

Using the same genome selection criteria and comparative genomic pipeline, a parallel analysis was performed for members of the *Bacteroidales* order. Representative *Bacteroidales* genomes were selected using the same quality threshold and retrieved from GTDB. Genome annotation, KEGG Ortholog assignment, KO presence-absence matrix construction, orthogroup inference, and species tree reconstruction were performed as described above, except that we used OrthoFinder v3.1.0^109^, running the analysis in two rounds (as suggested by its authors) for better scalability.

Trees were plotted using the R package ggtree^110^ v3.16.3 and gene neighborhoods were plotted using the R package gggenes^111^ v0.6.0.

### Reanalysis of metabolomics data

Metabolomics data were downloaded from the NIH Comon Fund’s National Metabolomics Data Repository (NMDR) website, the Metabolomics Workbench^112^ (https://www.metabolomicsworkbench.org). The specific study accession for Han et al. was ST001683 (project ID PR001074, doi:10.21228/M8W11P); the reversed phase positive ion mode “named metabolite” data were used. The Ho et al. study accession was ST002832 (project ID PR001774, available at doi:10.21228/M8DB1F) and the HILIC positive ion mode “named metabolite” data were used.

## Data availability

RNAseq data have been submitted to NCBI Gene Expression Omnibus (GEO) repository under accession number GSE319166. DNA sequencing data for the *R. peoriensis* genome is available from the NCBI Sequence Read Archive (SRA) under the BioProject PRJNA1076216.

## Acknowledgements

Funding for this project was provided by the NIH (R35GM151155 to PHB), by Ohio State University (startup funds to PHB), and by Jobs OHIO and Ohio State University (OSU Office of Enterprise, Research, Innovation, and Knowledge [ERIK] award to JAN). This study used data from the Metabolomics Workbench, which is supported by Metabolomics Workbench/National Metabolomics Data Repository (NMDR) (grant U2C-DK119886), the Common Fund Data Ecosystem (CFDE) (grant 3OT2OD030544) and the Metabolomics Consortium Coordinating Center (M3C) (grant 1U2C-DK119889). We also wish to acknowledge Arthur Franklin for helpful contributions.

## Supplementary Information

Supplementary Table S1: Detailed strain information for all strains used in this study.

Supplementary Table S2: Table of all KOfams present in *D. formicigenerans, R. peoriensis*, and *A. caccae* but not *C. scindens* and *B. coccoides*.

Supplementary Table S3: Table of all KOfams present in *C. scindens* and *B. coccoides* but not *D. formicigenerans, R. peoriensis*, or *A. caccae*.

Supplementary Table S4: *Lachnospiraceae* representative genomes used for calling orthogroups. Supplementary Table S5: Table of all potentially orthologous *fin* neighborhoods detected.

Supplementary Table S6: Bacteroidales representative genomes used for calling orthogroups.

Supplementary Figure S1: Read coverage plots across the *fin* neighborhood. A) 17.5µM riboflavin, 11 hours. B) 0.13µM riboflavin, 12 hours.

Supplementary Figure S2: Orthogroup gene trees for three genes in the fin neighborhood: A) bifunctional riboflavin kinase/FAD synthetase (*finB*), B) aldolase (*finC*), and C) aldehyde dehydrogenase (*finG*). The *C. scindens* and *F*.

*fissicatena* orthologs are labeled, and the clades selected for identifying potential orthologous neighborhoods are highlighted in yellow.

Supplementary Figure S3: Diagram showing detected *fin* gene neighborhoods in *C. scindens, M. intestini, F. fissicatena*, and *M. hominis*. Genes in the same orthogroup as a *C. scindens fin* gene are in pink; flanking genes that had a Prokka annotation (other than “hypothetical gene”) are labeled with their inferred function.

## References

1. Topping, D. L. & Clifton, P. M. Short-chain fatty acids and human colonic function: roles of resistant starch and nonstarch polysaccharides. Physiol. Rev. 81, 1031–1064 (2001).

2. Smith, P. M. et al. The microbial metabolites, short chain fatty acids, regulate colonic Treg cell homeostasis. Science 341, 10.1126/science.1241165 (2013).

3. Buffie, C. G. et al. Precision microbiome reconstitution restores bile acid mediated resistance to Clostridium difficile. Nature 517, 205–208 (2015).

4. Kang, J. D. et al. Bile Acid 7α-Dehydroxylating Gut Bacteria Secrete Antibiotics that Inhibit Clostridium difficile: Role of Secondary Bile Acids. Cell Chem. Biol. 26, 27-34.e4 (2019).

5. Aronov, P. A. et al. Colonic Contribution to Uremic Solutes. J. Am. Soc. Nephrol. 22, 1769 (2011).

6. Koeth, R. a et al. Intestinal microbiota metabolism of l-carnitine, a nutrient in red meat, promotes atherosclerosis. Nat. Med. 19, (2013).

7. Mastrangelo, A. et al. Imidazole propionate is a driver and therapeutic target in atherosclerosis. Nature 645, 254–261 (2025).

8. Roberts, A. B. et al. Development of a gut microbe–targeted nonlethal therapeutic to inhibit thrombosis potential. Nat. Med. 24, 1407–1417 (2018).

9. Gibson, G. R. & Roberfroid, M. B. Dietary modulation of the human colonic microbiota: introducing the concept of prebiotics. J. Nutr. 125, 1401–1412 (1995).

10. Zeng, X. et al. Gut bacterial nutrient preferences quantified in vivo. Cell 185, 3441-3456.e19 (2022).

11. Salyers, A. A., Vercellotti, J. R., West, S. E. & Wilkins, T. D. Fermentation of mucin and plant polysaccharides by strains of Bacteroides from the human colon. Appl. Environ. Microbiol. 33, 319–322 (1977).

12. Derrien, M. et al. Mucin-bacterial interactions in the human oral cavity and digestive tract. Gut Microbes 1, 254–268 (2010).

13. Culp, E. J. & Goodman, A. L. Cross-feeding in the gut microbiome: Ecology and Mechanisms. Cell Host Microbe 31, 485–499 (2023).

14. Basile, E. J., Shukla, K., Launico, M. V. & Sheer, A. J. Physiology, Nutrient Absorption. in StatPearls [Internet] (StatPearls Publishing, 2025).

15. Guillen, M. N., Rosener, B., Sayin, S. & Mitchell, A. Assembling stable syntrophic Escherichia coli communities by comprehensively identifying beneficiaries of secreted goods. Cell Syst. 12, 1064-1078.e7 (2021).

16. Magnúsdóttir, S., Ravcheev, D., de Crécy-Lagard, V. & Thiele, I. Systematic genome assessment of B-vitamin biosynthesis suggests co-operation among gut microbes. Front. Genet. 6, (2015).

17. Fenn, K. et al. Quinones are growth factors for the human gut microbiota. Microbiome 5, 161 (2017).

18. Soto-Martin, E. C. et al. Vitamin Biosynthesis by Human Gut Butyrate-Producing Bacteria and Cross-Feeding in Synthetic Microbial Communities. mBio 11, 10.1128/mbio.00886-20 (2020).

19. Belzer, C. et al. Microbial Metabolic Networks at the Mucus Layer Lead to Diet-Independent Butyrate and Vitamin B12 Production by Intestinal Symbionts. mBio 8, e00770–17 (2017).

20. Liu, L. et al. Riboflavin Supplementation Promotes Butyrate Production in the Absence of Gross Compositional Changes in the Gut Microbiota. Antioxid. Redox Signal. 38, 282–297 (2023).

21. Morrison, D. J. & Preston, T. Formation of short chain fatty acids by the gut microbiota and their impact on human metabolism. Gut Microbes 7, 189–200 (2016).

22. Fischbach, M. A. & Sonnenburg, J. L. Eating For Two: How Metabolism Establishes Interspecies Interactions in the Gut. Cell Host Microbe 10, 336–347 (2011).

23. Campbell, A., Gdanetz, K., Schmidt, A. W. & Schmidt, T. M. H2 generated by fermentation in the human gut microbiome influences metabolism and competitive fitness of gut butyrate producers. Microbiome 11, 133 (2023).

24. Kees, E. D. et al. Secreted Flavin Cofactors for Anaerobic Respiration of Fumarate and Urocanate by Shewanella oneidensis: Cost and Role. Appl. Environ. Microbiol. 85, e00852 (2019).

25. Kotloski, N. J. & Gralnick, J. A. Flavin electron shuttles dominate extracellular electron transfer by Shewanella oneidensis. mBio 4, e00553–12 (2013).

26. Light, S. H. et al. A flavin-based extracellular electron transfer mechanism in diverse Gram-positive bacteria. Nature 562, 140–144 (2018).

27. Khan, M. T. et al. The gut anaerobe Faecalibacterium prausnitzii uses an extracellular electron shuttle to grow at oxic–anoxic interphases. ISME J. 6, 1578–1585 (2012).

28. Dietrich, L. E. P., Teal, T. K., Price-Whelan, A. & Newman, D. K. Redox-active antibiotics control gene expression and community behavior in divergent bacteria. Science 321, 1203–1206 (2008).

29. Wang, Y., Kern, S. E. & Newman, D. K. Endogenous Phenazine Antibiotics Promote Anaerobic Survival of Pseudomonas aeruginosa via Extracellular Electron Transfer. J. Bacteriol. 192, 365–369 (2010).

30. Degnan, P. H., Taga, M. E. & Goodman, A. L. Vitamin B12 as a modulator of gut microbial ecology. Cell Metab. 20, 769–778 (2014).

31. Mok, K. C. et al. Identification of a Novel Cobamide Remodeling Enzyme in the Beneficial Human Gut Bacterium Akkermansia muciniphila. mBio 11, e02507–20 (2020).

32. Lerma-Ortiz, C. et al. ‘Nothing of chemistry disappears in biology’: the Top 30 damage-prone endogenous metabolites. Biochem. Soc. Trans. 44, 961–971 (2016).

33. Alhapel, A. et al. Molecular and functional analysis of nicotinate catabolism in Eubacterium barkeri. Proc. Natl. Acad. Sci. U. S. A. 103, 12341–12346 (2006).

34. Xu, H. et al. Identification of the First Riboflavin Catabolic Gene Cluster Isolated from Microbacterium maritypicum G10. J. Biol. Chem. 291, 23506–23515 (2016).

35. Kanazawa, H. et al. Two-Component Flavin-Dependent Riboflavin Monooxygenase Degrades Riboflavin in Devosia riboflavina. J. Bacteriol. 200, e00022–18 (2018).

36. West, D. W. & Owen, E. C. The urinary excretion of metabolites of riboflavine by man. Br. J. Nutr. 23, 889–898 (1969).

37. Chastain, J. L. & Mccormick, D. B. Clarification and Quantitation of Primary (Tissue) and Secondary (Microbial) Catabolites of Riboflavin That Are Excreted in Mammalian (Rat) Urine. J. Nutr. 117, 468–475 (1987).

38. Owen, E. C., West, D. W. & Coates, M. E. Metabolism of riboflavine in germ-free and conventional rabbits. Br. J. Nutr. 24, 259–267 (1970).

39. Foster, J. W. Microbiological Aspects of Riboflavin: I. Introduction. II. Bacterial Oxidation of Riboflavin to Lumichrome. J. Bacteriol. 47, 27–41 (1944).

40. Abdill, R. J. et al. Integration of 168,000 samples reveals global patterns of the human gut microbiome. Cell 188, 1100-1118.e17 (2025).

41. Devendran, S. et al. Clostridium scindens ATCC 35704: Integration of Nutritional Requirements, the Complete Genome Sequence, and Global Transcriptional Responses to Bile Acids. Appl. Environ. Microbiol. 85, e00052–19 (2019).

42. Kang, J. D. et al. Bile Acid 7α-Dehydroxylating Gut Bacteria Secrete Antibiotics that Inhibit Clostridium difficile: Role of Secondary Bile Acids. Cell Chem. Biol. 26, 27-34.e4 (2019).

43. Abdugheni, R. et al. Metabolite profiling of human-originated Lachnospiraceae at the strain level. iMeta 1, e58 (2022).

44. Noecker, C. et al. Systems biology elucidates the distinctive metabolic niche filled by the human gut microbe Eggerthella lenta. PLOS Biol. 21, e3002125 (2023).

45. Bisanz, J. E. et al. A Genomic Toolkit for the Mechanistic Dissection of Intractable Human Gut Bacteria. Cell Host Microbe 27, 1001-1013.e9 (2020).

46. Kananen, K. et al. Adaptive adjustment of profile HMM significance thresholds improves functional and metabolic insights into microbial genomes. Bioinforma. Adv. 5, vbaf039 (2025).

47. Meibom, K. L. et al. BaiJ and BaiB are key enzymes in the chenodeoxycholic acid 7α-dehydroxylation pathway in the gut microbe Clostridium scindens ATCC 35704. Gut Microbes 16, 2323233 (2024).

48. Harris, S. C. et al. Identification of a gene encoding a flavoprotein involved in bile acid metabolism by the human gut bacterium Clostridium scindens ATCC 35704. Biochim. Biophys. Acta BBA - Mol. Cell Biol. Lipids 1863, 276–283 (2018).

49. Buckel, W., Ermler, U., Vonck, J., Fritz, G. & Steuber, J. The RNF/NQR redox pumps: a versatile system for energy transduction in bacteria and archaea. Appl. Microbiol. Biotechnol. 109, 148 (2025).

50. Wilson, A. C. & Pardee, A. B. Regulation of Flavin Synthesis by Escherichia coli. Microbiology 28, 283–303 (1962).

51. Bacher, A., Eberhardt, S., Fischer, M., Kis, K. & Richter, G. Biosynthesis of vitamin b2 (riboflavin). Annu. Rev. Nutr. 20, 153–167 (2000).

52. Gutiérrez-Preciado, A. et al. Extensive Identification of Bacterial Riboflavin Transporters and Their Distribution across Bacterial Species. PLoS ONE 10, e0126124 (2015).

53. Burgess, C. M. et al. The Riboflavin Transporter RibU in Lactococcus lactis: Molecular Characterization of Gene Expression and the Transport Mechanism. J. Bacteriol. 188, 2752–2760 (2006).

54. Gelfand, M. S., Mironov, A. A., Jomantas, J., Kozlov, Y. I. & Perumov, D. A. A conserved RNA structure element involved in the regulation of bacterial riboflavin synthesis genes. Trends Genet. 15, 439–442 (1999).

55. Mironov, A. S. et al. Sensing Small Molecules by Nascent RNA: A Mechanism to Control Transcription in Bacteria. Cell 111, 747–756 (2002).

56. Haft, D. H. et al. RefSeq and the prokaryotic genome annotation pipeline in the age of metagenomes. Nucleic Acids Res. 52, D762–D769 (2023).

57. Hamada, K., Sasaki, M. & Yoshimura, K. Two New Types of Riboflavin-Decomposing Bacteria Isolated from Human Feces. J. Vitaminol. (Kyoto) 2, 307–315 (1956).

58. Taylor, M. M. Eubacterium fissicatena sp.nov. Isolated from the Alimentary Tract of the Goat. Microbiology 71, 457– 463 (1972).

59. Emms, D. M. & Kelly, S. OrthoFinder: phylogenetic orthology inference for comparative genomics. Genome Biol. 20, 238 (2019).

60. Uebanso, T., Shimohata, T., Mawatari, K. & Takahashi, A. Functional Roles of B-Vitamins in the Gut and Gut Microbiome. Mol. Nutr. Food Res. 64, 2000426 (2020).

61. U.S. Department of Agriculture, Agricultural Research Service & U.S. Department of Health and Human Services. What We Eat In America Data Tables, August 2021-2023. https://www.ars.usda.gov/northeast-area/beltsville-md-bhnrc/beltsville-human-nutrition-research-center/food-surveys-research-group/docs/wweia-data-tables/.

62. Zempleni, J., Galloway, J. & McCormick, D. Pharmacokinetics of orally and intravenously administered riboflavin in healthy humans. Am. J. Clin. Nutr. 63, 54–66 (1996).

63. Han, S. et al. A metabolomics pipeline for the mechanistic interrogation of the gut microbiome. Nature 595, 415–420 (2021).

64. Ho, P.-Y., Nguyen, T. H., Sanchez, J. M., DeFelice, B. C. & Huang, K. C. Resource competition predicts assembly of gut bacterial communities in vitro. Nat. Microbiol. 9, 1036–1048 (2024).

65. McAnulty, M. J. & Wood, T. K. YeeO from Escherichia coli exports flavins. Bioengineered 5, 386–392 (2014).

66. Oka, M. & McCormick, D. B. Urinary Lumichrome-Level Catabolites of Riboflavin are due to Microbial and Photochemical Events and not Rat Tissue Enzymatic Cleavage of the Ribityl Chain. J. Nutr. 115, 496–499 (1985).

67. Taylor, M. M. Eubacterium fissicatena sp.nov. Isolated from the Alimentary Tract of the Goat. J. Gen. Microbiol. 71, 457–463 (1972).

68. Reaves, M. L., Young, B. D., Hosios, A. M.Xu, Y.-F. & Rabinowitz, J. D. Pyrimidine homeostasis is accomplished by directed overflow metabolism. Nature 500, 237–241 (2013).

69. Spencer, M. E. & Guest, J. R. Isolation and properties of fumarate reductase mutants of Escherichia coli. J. Bacteriol. 114, 563–570 (1973).

70. Bogachev, A. V., Bertsova, Y. V., Bloch, D. A. & Verkhovsky, M. I. Urocanate reductase: identification of a novel anaerobic respiratory pathway in Shewanella oneidensis MR-1. Mol. Microbiol. 86, 1452–1463 (2012).

71. Harris, S. C. et al. Identification of a gene encoding a flavoprotein involved in bile acid metabolism by the human gut bacterium Clostridium scindens ATCC 35704. Biochim. Biophys. Acta Mol. Cell Biol. Lipids 1863, 276–283 (2018).

72. Farhana, A. et al. Reductive stress in microbes: implications for understanding Mycobacterium tuberculosis disease and persistence. Adv. Microb. Physiol. 57, 43–117 (2010).

73. Frelin, O. et al. A directed-overflow and damage-control N-glycosidase in riboflavin biosynthesis. Biochem. J. 466, 137–145 (2015).

74. Sebastián, M. et al. The FAD synthetase from the human pathogen Streptococcus pneumoniae: a bifunctional enzyme exhibiting activity-dependent redox requirements. Sci. Rep. 7, 7609 (2017).

75. Kearney, E. B., Goldenberg, J., Lipsick, J. & Perl, M. Flavokinase and FAD synthetase from Bacillus subtilis specific for reduced flavins. J. Biol. Chem. 254, 9551–9557 (1979).

76. Ahmad, R. & Armstrong, D. A. Reduction of Lumichrome by the Radical Anions of CO2 and Lipoamide. Int. J. Radiat. Biol. Relat. Stud. Phys. Chem. Med. 45, 607–614 (1984).

77. Bommer, G. T., Schaftingen, E. V. & Veiga-da-Cunha, M. Metabolite Repair Enzymes Control Metabolic Damage in Glycolysis. Trends Biochem. Sci. 45, 228–243 (2020).

78. Lerma-Ortiz, C. et al. ‘Nothing of chemistry disappears in biology’: the Top 30 damage-prone endogenous metabolites. Biochem. Soc. Trans. 44, 961–971 (2016).

79. Hartl, J., Kiefer, P., Meyer, F. & Vorholt, J. A. Longevity of major coenzymes allows minimal de novo synthesis in microorganisms. Nat. Microbiol. 2, 17073 (2017).

80. Meier, K. H. U. et al. Metabolic landscape of the male mouse gut identifies different niches determined by microbial activities. Nat. Metab. 5, 968–980 (2023).

81. Putnam, E. E. & Goodman, A. L. B vitamin acquisition by gut commensal bacteria. PLOS Pathog. 16, e1008208 (2020).

82. Rajamani, S. et al. The Vitamin Riboflavin and Its Derivative Lumichrome Activate the LasR Bacterial Quorum-Sensing Receptor. Mol. Plant-Microbe Interactions® 21, 1184–1192 (2008).

83. Gouws, L. M. et al. The Plant Growth Promoting Substance, Lumichrome, Mimics Starch, and Ethylene-Associated Symbiotic Responses in Lotus and Tomato Roots. Front. Plant Sci. 3, 120 (2012).

84. Sinha, S. et al. Vitamin B2 Catabolism: Nature’s Route from Riboflavin to Acetoacetate and Pyruvate. ACS Cent. Sci. 11, 2353–2365 (2025).

85. Kjer-Nielsen, L. et al. MR1 presents microbial vitamin B metabolites to MAIT cells. Nature 491, 717–723 (2012).

86. Corbett, A. J. et al. T-cell activation by transitory neo-antigens derived from distinct microbial pathways. Nature 509, 361–365 (2014).

87. Abdelaal, M. R. et al. The antigen-presenting molecule MR1 binds host-generated riboflavin catabolites. J. Exp. Med. 223, e20250711 (2025).

88. Wolin, E. A., Wolin, M. J. & Wolfe, R. S. Formation of Methane by Bacterial Extracts. J. Biol. Chem. 238, 2882–2886 (1963).

89. Sprouffske, K. & Wagner, A. Growthcurver: an R package for obtaining interpretable metrics from microbial growth curves. BMC Bioinformatics 17, 172 (2016).

90. Wick, R. R., Judd, L. M. & Holt, K. E. Assembling the perfect bacterial genome using Oxford Nanopore and Illumina sequencing. PLOS Comput. Biol. 19, e1010905 (2023).

91. Chen, S. fastp 1.0: An ultra‐fast all‐round tool for FASTQ data quality control and preprocessing. iMeta 4, e70078 (2025).

92. Wick, R. R. et al. Trycycler: consensus long-read assemblies for bacterial genomes. Genome Biol. 22, 266 (2021).

93. Kolmogorov, M., Yuan, J., Lin, Y. & Pevzner, P. A. Assembly of long, error-prone reads using repeat graphs. Nat. Biotechnol. 37, 540–546 (2019).

94. Wick, R. R. & Holt, K. E. Benchmarking of long-read assemblers for prokaryote whole genome sequencing. Preprint at 10.12688/f1000research.21782.4 (2021).

95. Vaser, R. & Šikić, M. Time- and memory-efficient genome assembly with Raven. Nat. Comput. Sci. 1, 332–336 (2021).

96. ananoporetech/medaka. Oxford Nanopore Technologies (2026).

97. Wick, R. R. & Holt, K. E. Polypolish: Short-read polishing of long-read bacterial genome assemblies. PLoS Comput. Biol. 18, e1009802 (2022).

98. Zimin, A. V. & Salzberg, S. L. The genome polishing tool POLCA makes fast and accurate corrections in genome assemblies. PLOS Comput. Biol. 16, e1007981 (2020).

99. Eren, A. M. et al. Community-led, integrated, reproducible multi-omics with anvi’o. Nat. Microbiol. 6, 3–6 (2021).

100. Bolger, A. M., Lohse, M. & Usadel, B. Trimmomatic: a flexible trimmer for Illumina sequence data. Bioinformatics 30, 2114–2120 (2014).

101. Love, M. I., Huber, W. & Anders, S. Moderated estimation of fold change and dispersion for RNA-seq data with DESeq2. Genome Biol. 15, 550 (2014).

102. Gu, Z. Complex heatmap visualization. iMeta 1, e43 (2022).

103. Seemann, T. Prokka: rapid prokaryotic genome annotation. Bioinformatics 30, 2068–2069 (2014).

104. Hyatt, D. et al. Prodigal: prokaryotic gene recognition and translation initiation site identification. BMC Bioinformatics 11, 119 (2010).

105. Price, M. N., Dehal, P. S. & Arkin, A. P. FastTree 2--approximately maximum-likelihood trees for large alignments. PloS One 5, e9490 (2010).

106. Katoh, K., Misawa, K., Kuma, K. & Miyata, T. MAFFT: a novel method for rapid multiple sequence alignment based on fast Fourier transform. Nucleic Acids Res. 30, 3059–3066 (2002).

107. Kelly, S. & Maini, P. K. DendroBLAST: approximate phylogenetic trees in the absence of multiple sequence alignments. PloS One 8, e58537 (2013).

108. Steinegger, M. & Söding, J. MMseqs2 enables sensitive protein sequence searching for the analysis of massive data sets. Nat. Biotechnol. 35, 1026–1028 (2017).

109. Emms, D. M., Liu, Y., Belcher, L., Holmes, J. & Kelly, S. OrthoFinder: scalable phylogenetic orthology inference for comparative genomics. 2025.07.15.664860 Preprint at 10.1101/2025.07.15.664860 (2025).

110. Yu, G., Smith, D., Zhu, H., Guan, Y. & Lam, T. T.-Y. ggtree: an R package for visualization and annotation of phylogenetic trees with their covariates and other associated data. Methods Ecol. Evol. 8, 28–36 (2017).

111. Wilkins, D. & Kurtz, Z. gggenes: Draw Gene Arrow Maps in ‘ggplot2’. (2025).

112. Sud, M. et al. Metabolomics Workbench: An international repository for metabolomics data and metadata, metabolite standards, protocols, tutorials and training, and analysis tools. Nucleic Acids Res. 44, D463–470 (2016).

